# Lipopeptide antibiotics disrupt interactions of undecaprenyl phosphate with UptA

**DOI:** 10.1101/2024.04.02.587717

**Authors:** Abraham O. Oluwole, Neha Kalmankar, Michela Guida, Jack L. Bennett, Giovanna Poce, Jani R. Bolla, Carol V. Robinson

## Abstract

The peptidoglycan pathway represents one of the most successful antibacterial targets with the last critical step being the flipping of carrier lipid, undecaprenyl phosphate (C_55_-P), across the membrane to re-enter the pathway. This translocation of C_55_-P is facilitated by DedA and DUF368 domain-containing family membrane proteins via unknown mechanisms. Here we employ native mass spectrometry to investigate the interactions of UptA, a member of the DedA family of membrane protein from *Bacillus subtilis*, with C_55_-P, membrane phospholipids and cell wall-targeting antibiotics. Our results show that UptA, expressed and purified in *E. coli*, forms monomer-dimer equilibria, and binds to C_55_-P in a pH-dependent fashion. Specifically, we show that UptA interacts more favourably with C_55_-P over shorter-chain analogues and membrane phospholipids. Moreover, we demonstrate that lipopeptide antibiotics, amphomycin and aspartocin D, can directly inhibit UptA function by out-competing the substrate for the protein binding, in addition to their propensity to form complex with free C_55_-P. Overall, this study shows that UptA-mediated translocation of C_55_-P is potentially mediated by pH and anionic phospholipids and provides insights for future development of antibiotics targeting carrier lipid recycling.

## Introduction

The global threat of drug-resistant bacterial pathogens calls for renewed efforts in the search for new antibiotics, along with the identification of new biochemical pathways and interactions that can be effectively inhibited. A key antibiotic target is peptidoglycan, the cell wall polymer that provides bacteria with resistance to osmotic stress and environmental assaults (1, 2). Accordingly, topline antibiotics such as amoxicillin and vancomycin (3, 4) inhibit the transpeptidation step of the peptidoglycan synthesis (5–7). Moreover, promising antibiotic candidates, such as ramoplanin and moenomycin, target the transglycosylation step (8, 9). Peptidoglycan biosynthesis begins with the formation of uridine diphosphate-*N*-acetylmuramyl-pentapeptide (UM5) in the cytosol (10), followed by its coupling to the lipid carrier undecaprenyl monophosphate (C_55_-P) on the cytosolic side of the cytoplasmic membrane (11, 12). The resulting lipid I is further decorated with a GlcNAc residue to form lipid II (13) before being flipped across the cytoplasmic membrane (14–16). In the periplasm, the headgroup of lipid II is polymerized and cross-linked into the existing meshwork (17–19). The lipid carrier is then released as undecaprenyl diphosphate (C_55_-PP). To participate in the next round of precursor transfer, C_55_-PP must be dephosphorylated to form C_55_-P (20–22) and then flipped across the cytoplasmic membrane such that its phosphate headgroup returns to the cytoplasmic side. The mechanism by which C_55_-P is translocated across the cytoplasmic membrane is the least understood of the membrane-associated steps of the peptidoglycan pathway (Fig. 1A).

**Fig. 1:**
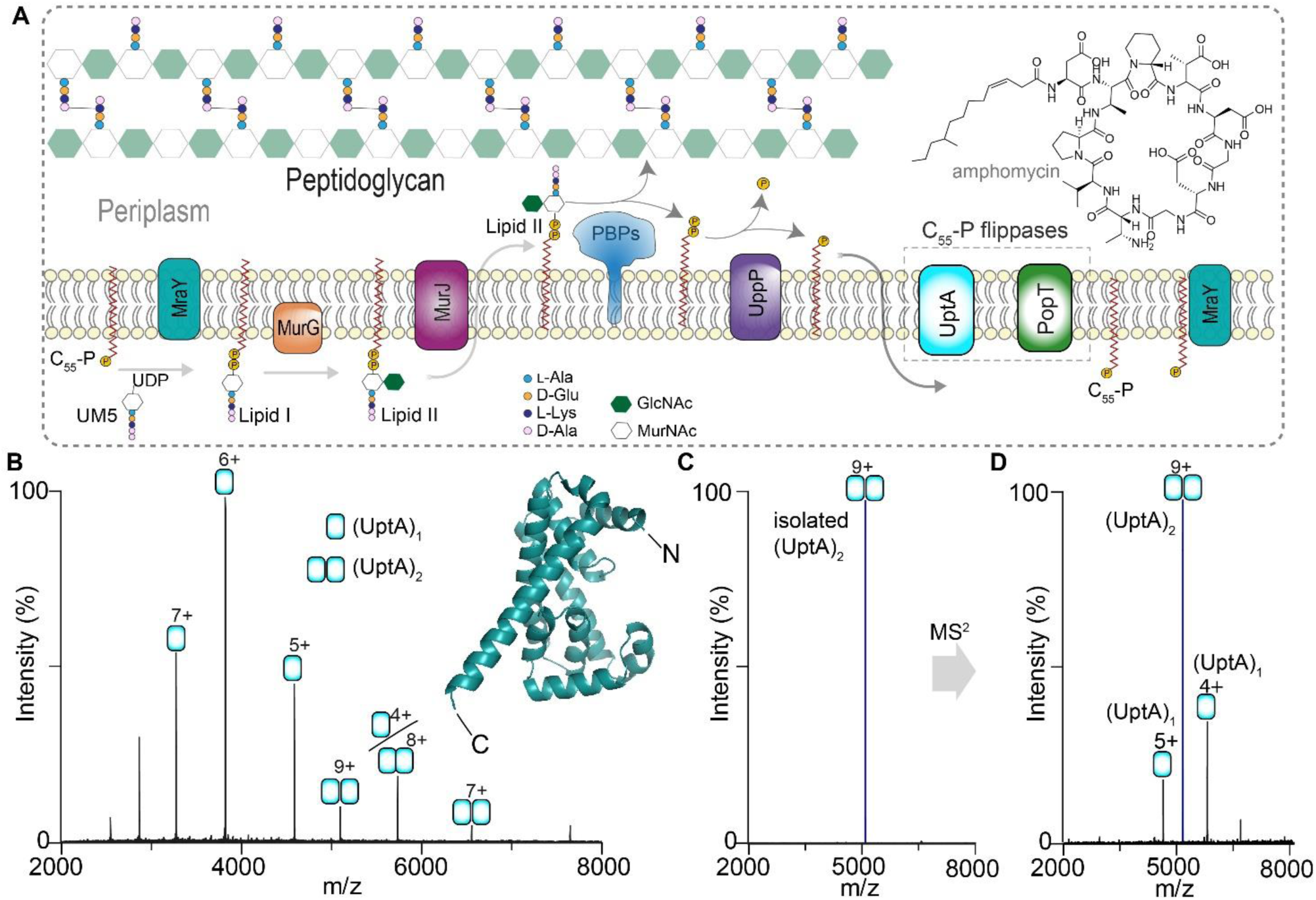
UptA is primarily monomeric with a low population of dimer. **A**) Schematic depiction of the lipid II cycle of peptidoglycan biosynthesis. C_55_-P, Undecaprenyl monophosphate; C_55_-PP, Undecaprenyl diphosphate; UM5, UDP-*N*-acetylmuramyl-pentapeptide; PBPs, penicillin-binding proteins. **B**) Native mass spectrum of UptA (2.2 µM) liberated from a buffer containing 200 mM ammonium acetate (pH 8.0), 0.05% LDAO. Peaks are assigned to UptA in monomeric and dimeric forms. Insert, the model structure of UptA (AFO31823-F1) predicted by AlphaFold (61). Predicted N- and C-termini are indicated. **C**) Quadrupole isolation of the UptA dimer (9+ charge state). **D**) MS/MS of the isolated UptA dimer upon collisional activation (190 V). The dimer dissociates into individual UptA protomers confirming their non-covalent association.

Recently, proteins belonging to the DedA- and the DUF368 domain-containing families were shown to facilitate trans-bilayer transport of C_55_-P in the Gram-positive bacteria *Bacillus subtilis* and *Staphylococcus aureus* (23), and in the Gram-negative bacteria *Vibrio cholerae* (24). DedA proteins are highly conserved, having 8 homologues in the model bacteria *E. coli* and 6 homologues in *B. subtilis* (25–27). Members of the DedA transporters in *B. subtilis* include UptA and PetA, which are proposed to facilitate trans-bilayer flipping of C_55_-P and phosphatidylethanolamine, respectively (23, 28). Compared to the DedA family, the DUF368-domain containing proteins, exemplified by PopT, are less conserved, being found in *S. aureus* and *V. cholerae,* but are absent in *B. subtilis* and *E. coli* (29). *V. cholerae* PopT becomes important for cell survival only under alkaline conditions (24). Several Gram-positive and Gram-negative bacteria encode one or multiple UptA-type transporters suggesting that the C_55_-P recycling process, mediated by these flippases, is broadly conserved across different bacterial species. For example, the single DedA homologue in *Borrelia burgdorferi* is an essential protein required for proper cell division (30). Therefore, the recycling step of peptidoglycan biosynthesis mediated by UptA-type flippases make promising antibiotics target (31). This necessitates a molecular-level understanding of how UptA interacts with its cognate substrate, C_55_-P, and potentially with membrane phospholipids, to elucidate the mechanism of C_55_-P translocation.

Herein we employ native mass spectrometry (native MS) to elucidate how *B. subtilis* UptA interacts with multiple lipidic substrates and antibiotics. We show that purified UptA exists as an equilibrium of monomers and dimers. We further show that UptA interacts more favorably with C_55_-P than C_55_-PP and membrane phospholipids, and then provide molecular-level evidence on the amino acid residues involved in C_55_-P binding. We find that UptA binds to its cognate ligand C_55_-P in a pH-dependent manner and show that lipopeptide antibiotics, such as amphomycin and aspartocin D, bind to UptA with higher affinity than other cell-wall targeting antimicrobial peptides such as bacitracin and vancomycin. Interestingly, amphomycin and its analogues out-compete the flippase UptA for C_55_-P binding, raising the possibility of a new mode of action and avenue for the development of novel antibiotics to inhibit peptidoglycan biosynthesis.

## Results

### Monomer-dimer equilibrium of UptA

We expressed *B. subtilis* UptA in *E. coli* and introduced the purified protein into the mass spectrometer from a buffer containing 0.05% LDAO and 200 mM ammonium acetate (pH 8.0). The mass spectrum of UptA displays two charge state distributions assigned to monomers (22963.49±0.70 Da) and dimers (45925.21±0.65 Da) (Fig. 1B). To confirm the non-covalent association of UptA monomers, we isolated the 9+ charge state assigned to the dimer and subjected these ions to high-energy collisional dissociation (HCD, Fig. 1C). The resulting MS^2^ spectrum displays new peaks that correspond by mass to UptA protomers, and bear charges complementary to the isolated parent ions (+5 and +4) (Fig. 1D). Of note, the experimental mass for the UptA monomer is higher than the theoretical sequence mass (22936.36 Da) by 28 Da, suggesting an endogenous modification. To identify this unknown modification, we activated the monomer (6+ charge state) using HCD, yielding a plethora of *b*- and *y-*type fragment ions (Fig. S1). The observed fragments are consistent with the formylation of the N-terminal methionine of UptA (Fig. S1). Bacterial inner membrane proteins are normally co-translationally formylated on the N-terminus, however, this modification is rarely observed *in vitro* due to subsequent removal by the cytosolic deformylase (32). Therefore, retainment of N-terminal formylation on UptA suggests periplasmic localisation of the N-terminus of UptA, thus supporting the predicted N-out topology (23). Additionally, we observe UptA in monomeric and dimeric forms in other detergents (Fig. S2), confirming the oligomeric composition of UptA, which might apply to other bacterial DedA homologues (33).

### UptA binds C_55_-P with high affinity

We explored the binding affinity of UptA towards C_55_-P by recording mass spectra for solutions containing 2.5 µM UptA and 0 to 40 µM C_55_-P in 200 mM ammonium acetate, 0.05% LDAO at pH 8.0. The mass spectrum for the equimolar mixture of UptA and C_55_-P yielded peaks corresponding to UptA in its apo form and in complex with C_55_-P (Fig. 2A), an indication of high-affinity binding interactions. Further increase in the concentration of C_55_-P enhanced the intensity of the UptA:C_55_-P complex, together with the emergence of additional binding events (Fig. 2A). We deconvoluted the spectra (34) to extract mean relative intensities of bound and unbound forms of UptA in the spectral series. The data reflected an increase in the relative abundance of UptA:(C_55_-P)_n_ complexes as a function of the concentration of C_55_-P (Fig. 2B). For the first binding event, the relative intensity plateaued at a protein/ligand molar ratio of ∼4 (Fig. 2B). By fitting the experimentally observed intensity ratios to the Hill equation, we obtained an apparent *K*_d_ = 5.7 µM (95% CI: 5.0 to 6.4 µM) for the binding of one C_55_-P molecule to UptA. This *K*_d_ value for UptA/C_55_-P binding is of the same order of magnitude as the binding interactions for MurG/lipid I (*K*_d_ = 1.89±0.6 µM) (35) and MurJ/lipid II (*K*_d_ = 2.9±0.6 µM) (14), indicating that UptA exhibits a similarly high affinity binding with respect to C_55_-P.

**Fig 2.**
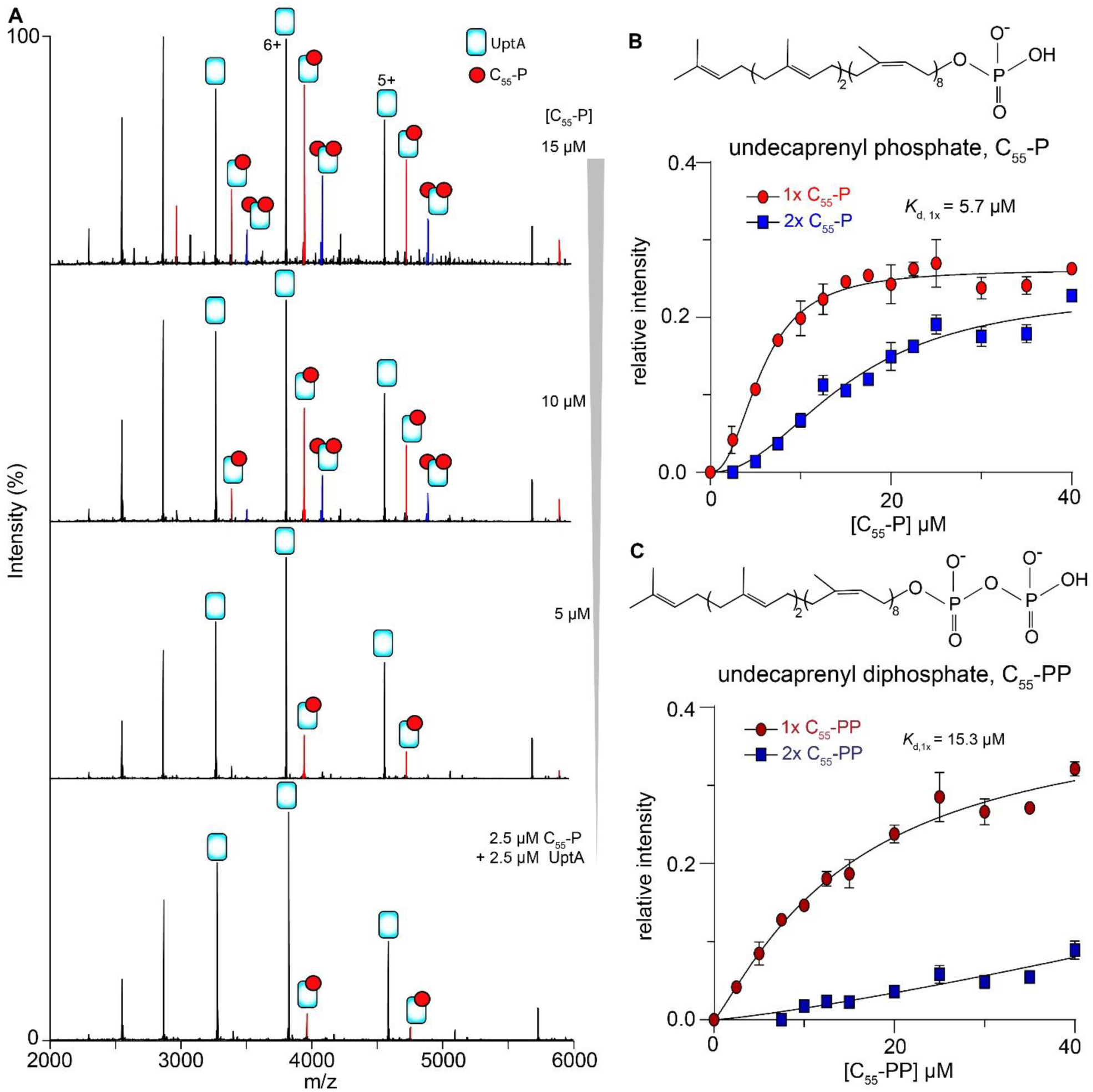
UptA binds C_55_-P with a high affinity. **A**) Spectra for UptA incubated with different concentrations of C_55_-P in a buffer containing 200 mM ammonium acetate (pH 8), 0.05% LDAO. **B-C**) The relative abundance of UptA bound to one (1x) or two (2x) molecules of C_55_-P (panel B) and C_55_-PP (panel C). Inserts, chemical structures of C_55_-P and C_55_-PP. Each data point represents an average of 3 replicate measurements, and the curves represent best fits of the data to Hill’s equation (see methods), error bars are standard deviations.

To investigate the binding selectivity of UptA for monophosphate (C_55_-P) versus diphosphate (C_55_-PP) forms of the lipid carrier, we performed comparative binding assays. First, we recorded mass spectra for a mixture of UptA with increasing concentrations of C_55_-PP (Fig. S3). We observed peaks assigned to UptA in ligand-free and C_55_-PP-bound forms, and the relative intensity of the UptA:C_55_-PP complex increased with increasing concentration of C_55_-PP (Fig. 2C). We then fit the data to Hill equation to obtain an apparent K_d_ for this system. The relative intensities of UptA:C_55_-PP yielded a significantly higher apparent K_d_ = 15.3 µM (95% CI: 9.5 to 32.8 µM) than the K_d_ obtained for UptA:C_55_-P (K_d_ = 5.7±0.7 µM). Being within an order of magnitude, these dissociation constants suggest that UptA forms extensive contact with both forms of the lipid carriers, but that the protein interacts more favourably with C_55_-P than C_55_-PP. We therefore conclude that UptA is more selective towards C_55_-P than C_55_-PP, thus confirming its preferred endogenous substrate.

### C_55_-P binding to UptA is sensitive to pH

UptA is a member of the DedA family of transporters whose cellular functions are potentially driven by proton-motive force and provide conditional fitness to bacteria under alkaline conditions (25, 26, 36). Membrane flippases such as MurJ, which is driven by proton-motive force (37), exhibit pH-dependency in ligand binding (14). We therefore considered the possibility that substrate interactions of UptA might exhibit pH dependency. To this end, we equilibrated C_55_-P with UptA at a protein/ligand molar ratio of 1:4 in a buffer of the same composition but at pH 8.0 and 5.0. The resulting spectrum at pH 8.0 exhibits peaks consistent with monomeric UptA in an apo form, and in complex with 1 and 2 molecules of C_55_-P (Fig. S4). We also observed UptA dimer in complex with up to two C_55_-P molecules, and the relative intensities of C_55_-P bound to the dimer is higher than to the monomer (Fig. S4). This suggested that dimer interacts more favourably with C_55_-P than the monomer UptA. However, the peaks assigned to ligand-bound UptA are more intense at pH 8.0 compared to pH 5.0 by a factor of 3. These results imply that more favourable protein-ligand interactions occur under physiologically relevant pH conditions. As a negative control, we tested for pH sensitivity of C_55_-P binding to MraY, the latter being the downstream enzyme in the peptidoglycan pathway that couples UM5 to C_55_-P. To this end, we equilibrated MraY and C_55_-P at a molar ratio of 1:4 in a buffer of pH 8.0 and pH 5.0. In this case, the spectra show no significant difference in the intensity of C_55_-P bound to MraY at pH 8.0 and at pH 5.0 (Fig. S4). We therefore conclude that environmental pH modulates the interaction of C_55_-P with UptA, and potentially its transport mechanism.

### Amino acid residues mediating C_55_-P interactions with UptA

Conserved arginine residues in the putative membrane re-entrant loops of *E. coli* DedA proteins YqjA (R130) and YghB (R136) are important for the cellular role of these proteins (36). The corresponding arginine residues (R112 and R118) in *B. subtilis* UptA have been suggested to impair the cellular function of UptA by making the cells more susceptible to inhibition by amphomycin (23). How these residues impact substrate interactions, however, is not known. Based on the model structure of UptA predicted by AlphaFold, the R112 residue should engage Q64 while R118 is able to form hydrogen bonds with E32, H119 and W146 to potentially stabilise the membrane re-entrant loops (Fig. 3A). To test for the impact of these residues on C_55_-P binding, we generated the E32A, Q64A, R112A, R118A, H119A and W146A UptA single mutants. After purification of the proteins, using the same protocol as the wild type (WT) (See Methods and Fig. S5), we equilibrated aliquots of each protein variant (5 µM) with C_55_-P (10 µM) and recorded spectra under the same instrument settings. The resulting spectra exhibited peaks consistent with C_55_-P binding in all cases; but only a modest reduction in C_55_-P binding intensity was observed in the case of the Q64A and H119A mutants compared to the wild type (Fig. S5, Fig. 3B). Compared to the wild type UptA, we observed more significant reduction (27-30%) in C_55_-P binding with the mutants E32A, W146A, R112A and R118A that participate directly in the predicted hydrogen bonding network (Figure 3A, B), suggesting that these residues are important for C_55_-P binding. We probe this observation further by investigating the double mutants R112A/R118A and R118A/R119A where the C_55_-P binding reduced more significantly (43% and 46%, respectively) than in the single mutants. We additionally generated a double mutant R112E/R118E in which the polarity of both native arginine residues in the wild type are reversed. We find that the R112E/R118E UptA exhibited 63% reduction in the intensity of C_55_-P binding compared to the wild type (Fig. 3B). These observations suggest that impaired hydrogen bonding between R118 and Q64 on the one hand, and R118 and E32 could have also disrupted the C_55_-P binding sites, highlighting the roles of R112 and R118 in the binding of C_55_-P to UptA.

**Fig. 3.**
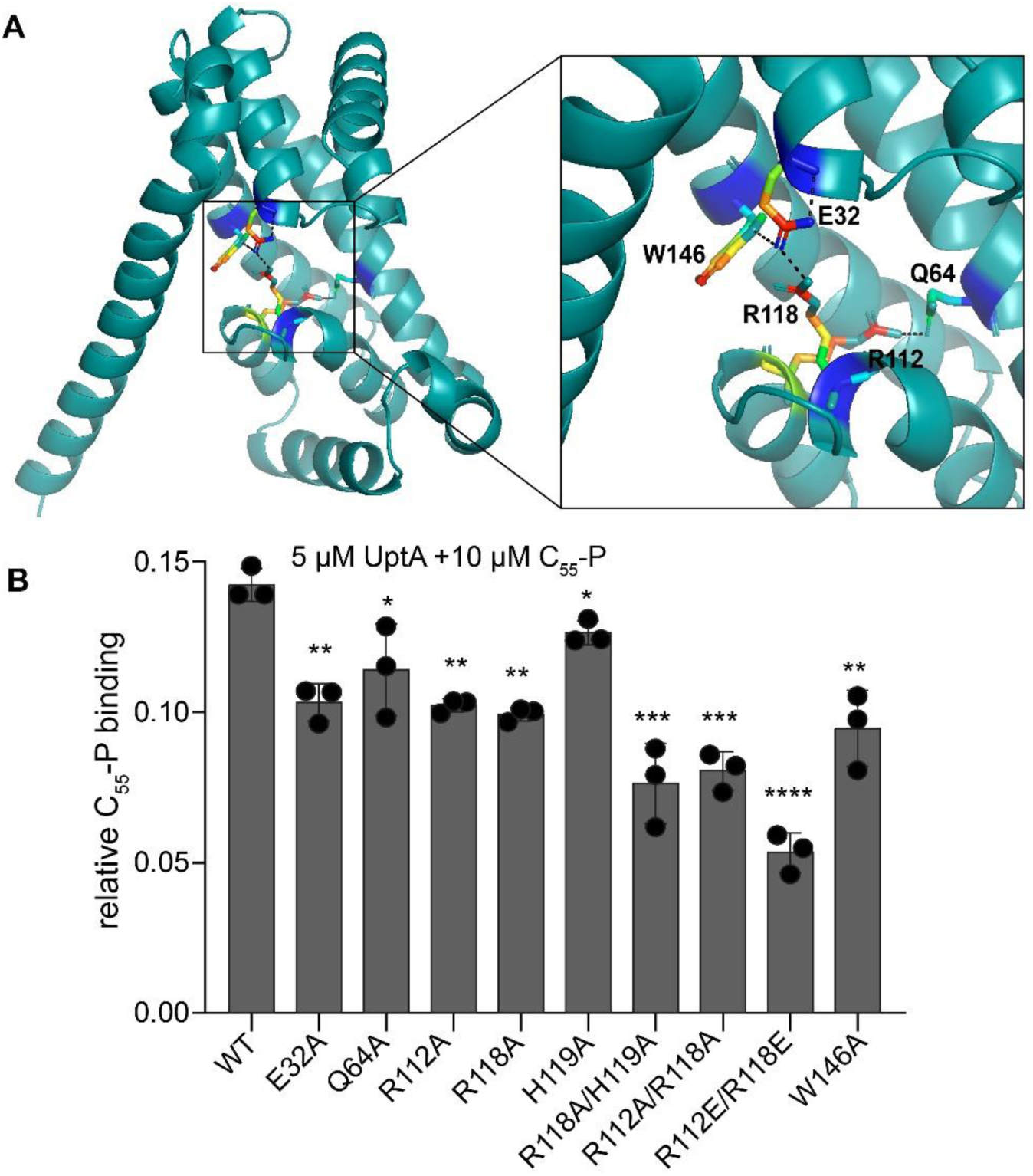
Probing the amino acid residues involved in UptA-C_55_-P binding. **A.** Model UptA structure predicted by AlphaFold (AFO31823-F1) (61) and a zoomed view showing hydrogen bonding networks surrounding R118 and R112. R112--Q64: 3.1 Å; R118--E32: 2.7 Å; and E32--W146: 3.5 Å. Hydrogen bonding distances were computed using ProteinTools(62) and the image was processed in PyMOL. **B.** Relative intensity of C_55_-P bound to UptA wild type and mutants. Bars represent the mean of three biological replicates with each data point shown in black, error bars are standard deviations. The double mutant at residues R112 and R118 caused the most significant reduction in C_55_-P binding. \**p* = 0.02 to 0.04, \*\**p* = 0.004 to 0.0012, \*\*\*\**p* = 0.0002, \*\*\*\**p* < 0.0001 from unpaired two-tailed t-test by comparing mutants with wild type, n = 3.

Since wild-type UptA displayed monomer-dimer equilibria and the dimer bind to C_55_-P more intensely than the monomer (cf. Fig. S4), we probed this observation further using the UptA single amino acid mutant E32A since the latter consistently retained a higher dimer population than the wild type (Fig. S6). We prepared and analysed solutions containing 2.5 µM E32A UptA and 10 µM C_55_-P. The resulting spectra exhibited peaks assigned to UptA monomers and dimers in complex with C_55_-P (Fig. S6). We find that the relative intensity of ligand-bound dimers is significantly higher than the ligand-bound monomers, and the ligand binding has only a modest impact on the monomer-dimer ratios. Together these results suggest that the ligand C_55_-P binds to dimeric UptA with a higher affinity than to the monomeric form.

### UptA interacts more favourably with C_55_-P than its shorter-chain analogues

Next, we investigated the chain length selectivity of UptA by testing its interaction with geranylgeranyl phosphate (C_20_-P), hexaprenyl phosphate (C_30_-P), and C_55_-P; the latter representing the native form of the lipid carrier in the model organisms *B. subtilis* and *E. coli* (38). For this experiment, we incubated 4 µM UptA with a 10-fold molar excess of C_20_-P, C_30_-P and C_55_-P at pH 8.0 and recorded spectra under the same conditions. For the UptA/C_20_-P sample, the spectrum exhibits low-intensity peaks corresponding by mass to UptA:C_20_-P complexes, leaving most of the protein in the apo form (Fig. S7). In the case of UptA/C_30_-P, we observed more intense protein/ligand complexes, indicating a higher binding affinity of UptA to C_30_-P compared to C_20_-P (Fig. S7). The spectra recorded for the UptA/C_55_-P mixture present peaks with the highest intensity of protein/ligand complexes (Fig. S7), indicating a more favourable interaction for C_55_-P compared to the lipidic substrate with shorter aliphatic chains. The length and stereochemistry of lipid tails can influence their interactions with membrane proteins (39). Our data shows that the lipid carrier with longer hydrophobic tails binds more favourably to UptA than the shorter ones, supporting our hypothesis that C_55_-P can form extensive contact with UptA to favour its binding interactions. Accordingly, the relative abundance of UptA in complex with C_55_-P is higher than in the case of UptA-bound C_30_-P and C_20_-P (Fig. S7). Together, these data are consistent with C_55_-P being the preferred endogenous substrate for UptA.

### UptA binds more favourably to phosphatidylglycerols than phosphatidylethanolamines

Proteins belonging to the DedA family are a widespread family of transporters and are proposed to flip diverse lipids, including carrier lipids and membrane phospholipids. We therefore probed the interaction of UptA with membrane phospholipids to understand if the latter could modulate its function. For this purpose, we selected anionic phosphatidylglycerols (PG) and zwitterionic phosphatidylethanolamines (PE), representing the major classes of phospholipid headgroups found in *B. subtilis* (40, 41). We recorded mass spectra of solutions containing 4 µM UptA incubated with 10 µM a15:0-i15:0 PE and 10 µM a15:0-i15:0 PG (a15:0, anteiso-pentadecanoic acid; i15:0, iso-pentadecanoic acid). The fatty acid composition of these lipids closely mimics those of native *B. subtilis* lipids (see below). The resulting spectra display peaks corresponding to UptA in apo and lipid-bound forms in both cases (Fig. 4A,B). We observed the formation of 1:1 and 1:2 protein-lipid complexes of UptA with a15:0-i15:0 PG, but only a 1:1 complex in the case of a15:0-i15:0 PE. Accordingly, the mean relative intensity of UptA in complex with a15:0-i15:0 PG is higher than for a15:0-i15:0 PE (Fig. 4C). We therefore hypothesise that UptA interacts more favourably with PG than with PE. We tested additional phospholipids PG and PE with comparable acyl chains. We find that 16:0-18:1 PG and 18:1-18:1 PG bind to UptA more intensely than 16:0-18:1 PE and 18:1-18:1 PE, respectively (Fig. 4C), supporting the hypothesis. Importantly, UptA binds to C_55_-P more intensely than any of these diacylglycerol phospholipids (Fig. 4C), in line with C_55_-P being its native substrate.

**Fig 4.**
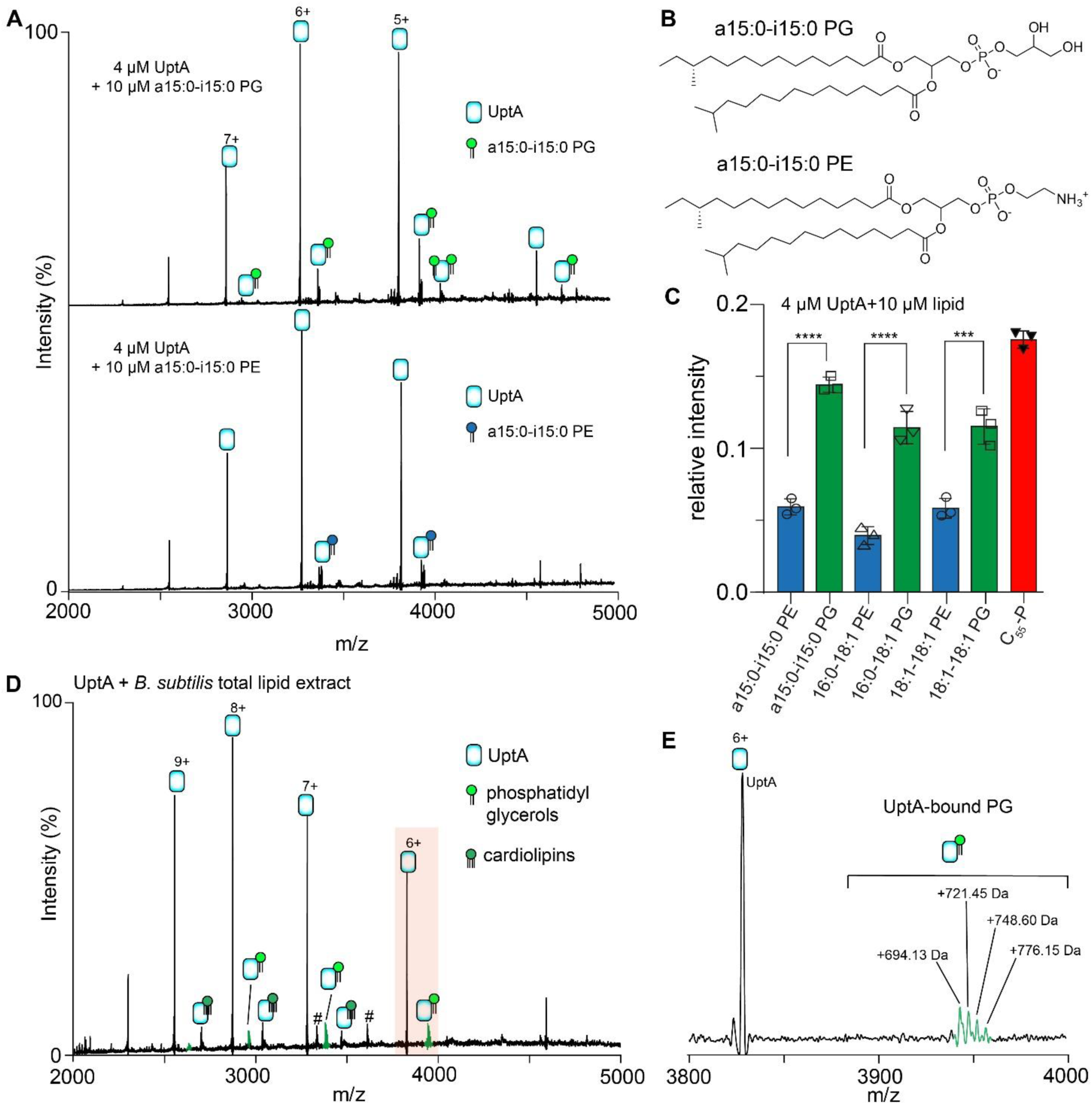
UptA binds phosphatidylglycerols. A) Native mass spectrum for UptA equilibrated with a15:0-i15:0 PE and a15:0-i15:0 PG. UptA interacts more favourably with PG than PE. **B)** Chemical structure of a15:0-i15:0 PE and a15:0-i15:0 PG. **C)** Relative intensity of UptA bound to phospholipids and C_55_-P. UptA interacts more favourably with C_55_-P than with the phospholipids. Bar represents the mean of three independent replicates shown as data points and the error bars are standard deviations. *p*-values are from two-tailed t-tests with n=3, \*\*\**p*=0.0003 to 0.0005, \*\*\*\**p*<0.0001. D. Native mass spectrum of 4 µM equilibrated with ∼0.1 mg/mL total lipid extracts of *B. subtilis.* Peaks labelled # is a 43.3 kDa contaminant. **E**. Zoomed view of a change state (6+), showing UptA adducts with phosphatidylglycerols.

To verify the lipid binding preference of UptA with more native-like lipids, we prepared lipid extracts from *B. subtilis* membranes (see Methods). We then incubate an aliquot of the resulting crude lipids (Fig. S8) with UptA. The spectra yielded peaks assigned to UptA in complex with cardiolipins (∼1326 Da) and a range of phospholipids, predominantly 694 Da, 721 Da, 748 Da and 776 Da species (Fig. 4D, 4E). To identify these lipids, we performed tandem MS/MS, isolating the lipid species and subjecting it to collisional activation. The resulting spectra exhibited fragments at 152.99 Da (Fig. S9), a diagnostic feature of the phosphatidylglycerol headgroup (42). The fatty acid fragments are mixed populations, predominantly 15:0-15:0, 15:0-17:0, 16:0-16:0, and 16:0-18:1 (Fig. S9) which are the typical fatty acids from *B. subtilis* membrane phospholipids (41). We therefore conclude that UptA interacts more favourably with the anionic cardiolipins and phosphatidylglycerols than with the zwitterionic phosphatidylethanolamines in the membrane lipid extracts.

### C_55_-P outcompetes phospholipids for UptA binding

To investigate whether or not C_55_-P can outcompete phospholipids in binding to UptA, we incubated solutions containing UptA-DOPG with increasing concentrations of C_55_-P. Spectra were recorded after aliquots of the solution were equilibrated on ice for 15 mins. The spectrum of UptA/DOPG mixture in the absence of C_55_-P reflected the binding of 1-2 molecules of DOPG per UptA (Fig. S10). The presence of C_55_-P in the sample yielded a new peak corresponding to UptA:C_55_-P complex, and simultaneously caused attenuation of the intensity DOPG-bound UptA in a concentration-dependent manner. At the highest C_55_-P concentration tested (20 µM), C_55_-P has displaced most of DOPG from UptA (Fig. S10). We therefore hypothesised that phospholipids should be unable to outcompete C_55_-P bound to UptA. To test this hypothesis, we incubated UptA with C_55_-P and then equilibrated aliquots with an increasing concentration of DOPG. Indeed, C_55_-P remains bound to UptA even in the presence of excess DOPG (20 µM) (Fig. S10), indicating that phospholipids could not significantly displace C_55_-P from UptA. Analogous experiments with DOPE show that C_55_-P effectively outcompete DOPE for UptA binding but the latter could not efficiently displace C_55_-P from UptA (Fig. S10). Together this study shows that UptA interacts more favourably with C_55_-P than with membrane phospholipids, thereby establishing C_55_-P as the preferred endogenous substrate.

### Lipopeptide antibiotics disrupt UptA:C_55_-P complex

We next investigate how UptA and its complex with C_55_-P interacts with a range of cell-wall targeting antibiotics including the lipopeptides daptomycin and amphomycin. Daptomycin is known to kill bacteria by depolarising the membrane (43) and by complexing with lipid II (44). We investigated if daptomycin affects C_55_-P binding to UptA by recording spectra for solutions containing 2.5 µM UptA, 10 µM C_55_-P and different concentrations of daptomycin. In the absence of antibiotics, the spectrum exhibited peaks assigned to UptA in apo form, and in complex with 1 and 2 C_55_-P molecules (Fig. S11). In the presence of daptomycin, we observed additional peaks including those corresponding to the binary complex UptA:daptomycin and the ternary complex UptA:C_55_-P:daptomycin (Fig. S11). Further increases in daptomycin concentration resulted in higher intensity of these complexes, indicating that daptomycin interacts with UptA but it does not directly compete against C_55_-P binding to the protein.

Amphomycin and derivatives are known to kill bacteria by forming complexes with free C_55_-P (45), however, whether or not this involves a direct interaction with a protein cellular target is not known. We therefore investigated how amphomycin affects the UptA:C_55_-P complex. We recorded spectra for solutions containing 2.5 µM UptA, 10 µM C_55_-P and then equilibrated aliquots with increasing concentrations of amphomycin. The spectra exhibited peaks consistent with amphomycin binding to UptA, in addition to the C_55_-P binding (Fig. 5A). We find that the peak intensities of the UptA:C_55_-P complex reduce with respect to an increase in the concentration of amphomycin (Fig. 5B). Compared to the case of daptomycin, little-to-no UptA:C_55_-P complexes remained in the presence of 10-fold molar excess of amphomycin (Fig. 5B), suggesting that amphomycin destabilise the ternary complex to sequester C_55_-P. We then performed similar experiments with aspartocin D, an analogue of amphomycin. The resulting spectrum shows that aspartocin D similarly binds to UptA and also induces dissociation of UptA:C_55_-P complex (Fig. 5C).

**Fig. 5.**
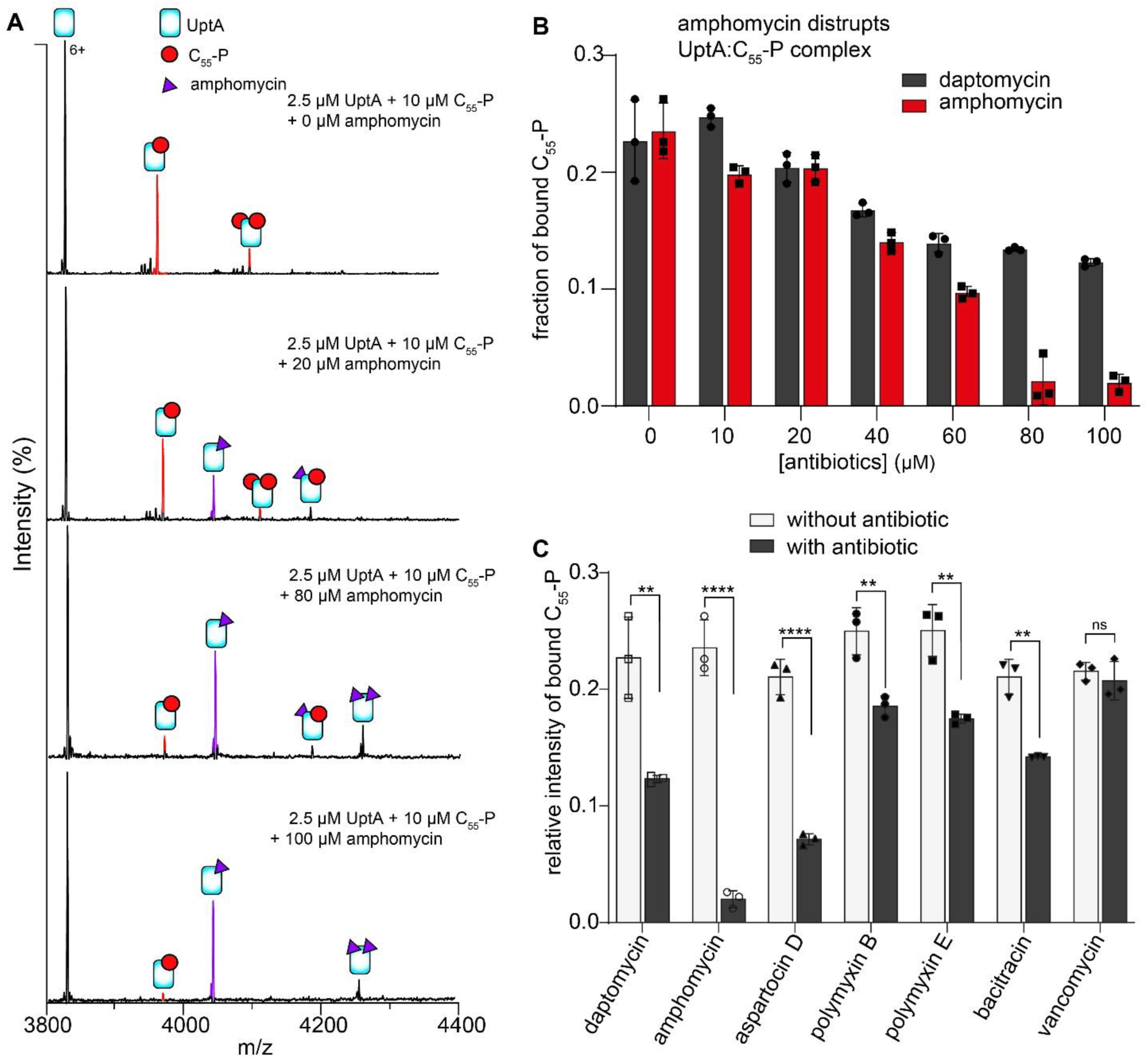
UptA function can be inhibited by lipopeptide antibiotics. **A)** Mass spectra (6+ charge state) for solutions containing 2.5 µM UptA and 10 µM C_55_-P incubated with 0-100 µM amphomycin. **B**) Relative intensity of C_55_-P bound to UptA in the presence of amphomycin and daptomycin. Bar represents the mean of three independent replicates shown as data points and error bars are standard deviations. Amphomycin destabilises the UptA:C_55_-P complex whereas daptomycin has a modest impact. **C)** Fraction of C_55_-P bound to UptA in the presence of cell-wall targeting antibiotics. Each sample contains 2.5 µM UptA, 10 µM C_55_-P and 100 µM of indicated antibiotic, and spectra were acquired using the same instrument settings and collisional activation of 75 V. Bar represents the mean of three independent replicates shown as data points, and error bar represents standard deviations. P-values from two-tailed t-tests with n=3; \*\**p*=0.001 to 0.006; ****p<0.0001, ns, not significant.

The amphiphilic nature of daptomycin and amphomycin raises the possibility that these lipopeptide antibiotics may promiscuously bind to UptA in a nonspecific manner. We probe this possibility by recording spectra for UptA:C_55_-P solutions in the presence of 100 µM polymyxin B and 100 µM polymyxin E. These polymyxins are membrane-disrupting lipopeptides that target the outer membrane lipopolysaccharides in Gram-negative bacteria (46) but without a known affinity for C_55_-P or UptA. We find that, unlike daptomycin and amphomycins, the polymyxins do not bind to UptA (Fig. S11), indicating that not every lipopeptide can bind to UptA. The ability of daptomycin and amphomycin analogues to directly interact with UptA therefore suggests that these lipopeptides may render a fraction of the cellular pool of UptA inaccessible to C_55_-P. Notably, polymyxin B and polymyxin E similarly caused attenuation of C_55_-P binding to UptA (Fig. S10), but this effect is less significant compared to the impact of amphomycin analogues (Fig. 5C). The reduction of C_55_-P binding to UptA in the presence of polymyxins can be attributed to potential electrostatic interactions between the phosphate group of C_55-_P molecules and the polymyxins which are cationic at physiological pH (47).

To verify these results, we studied the impact of bacitracin and vancomycin on UptA:C_55_-P interactions. Bacitracin and vancomycin are peptide antibiotics whose mode of action involves sequestration of C_55_-PP and lipid II, respectively (48, 49). To this end, we prepared UptA:C_55_-P mixtures and then equilibrated aliquots with bacitracin and vancomycin. We find that bacitracin and vancomycin bind to UptA but with very low intensities (Fig. S11), indicating weak affinity interactions. In line with the specificity of bacitracin for C_55_-PP rather than C_55_-P, this antibiotic modestly competes against C_55_-P binding to UptA (Fig. S11, Fig. 5C). By contrast, vancomycin has no observable impact on C_55_-P binding (Fig. S11, Fig. 5C). Thus, the non-lipopeptide antibiotics exhibit weak affinity for UptA and had only a minor impact on UptA:C_55_-P interactions. Taken together, our results highlight that the lipopeptides such as amphomycin and aspartocin D induce dissociation of UptA:C_55_-P complexes, and may therefore interfere with peptidoglycan pathway by binding directly to UptA in addition to their known propensity to form complexes with free C_55_-P molecules (45).

## Discussions

In this study, using native MS, we investigated the interplay of multiple lipidic substrates and antibiotics with UptA (summarised in Fig. 6). Specifically, we report that purified UptA exists as an equilibrium of monomers and dimers, suggesting that DedA proteins might form dimers in the native membrane when lateral pressures in the lipid bilayers (50) might prevent their dissociation. In line with the dependency of DedA family proteins on proton motive force (51), we observed less C_55_-P binding to UptA in the presence of excess H^+^ and that the monomer-dimer ratio of UptA is sensitive to pH rather than to C_55_-P binding. This feature contrasts with those of the upstream peptidoglycan enzymes MraY (35, 52) and UppP (53, 54) whose dimerization are promoted by lipid binding.

**Fig. 6.**
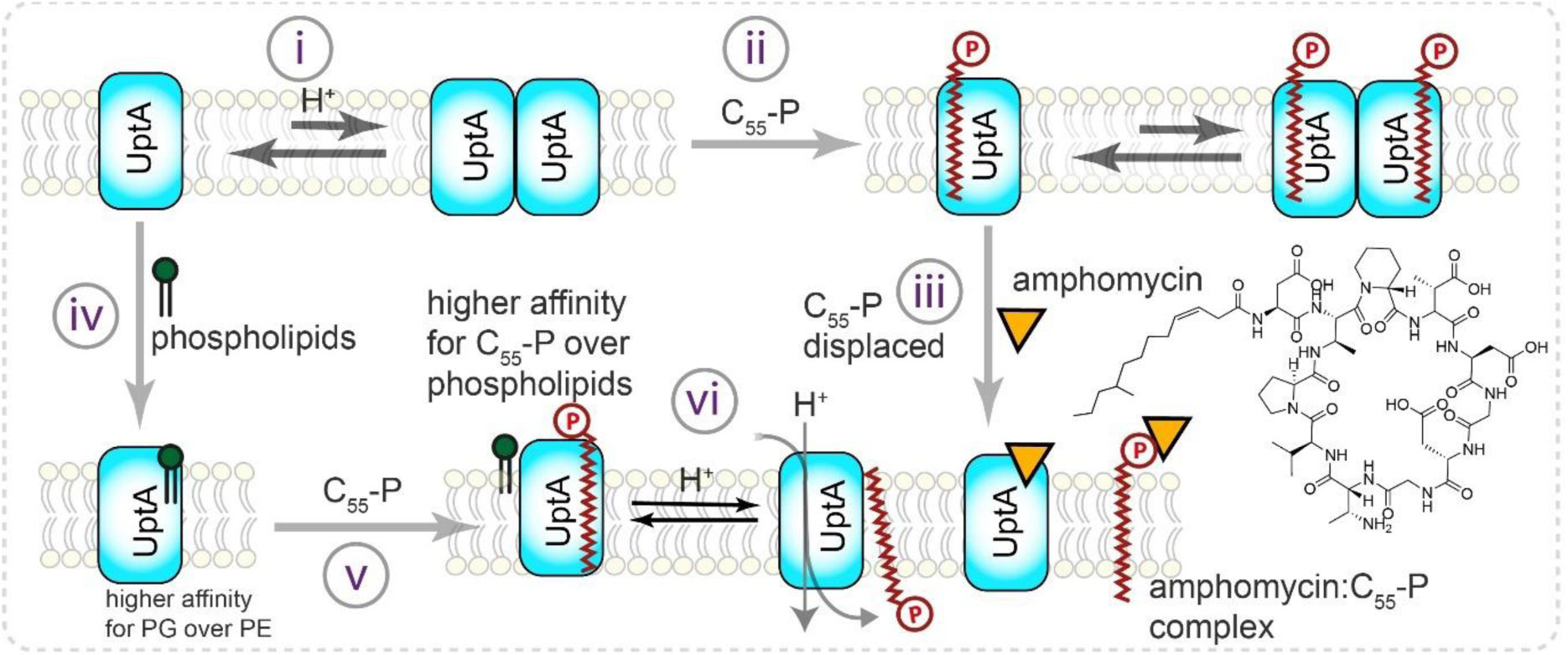
Insights from native MS into the interaction of UptA with lipidic substrates and antibiotics. (i) UptA exhibits monomer-dimer equilibria, with more dimer at pH 8.0 than pH 5.0. (ii) UptA interacts with C_55_-P more favourably than C_55_-PP, C_30_-P, and C_20_-P. (iii) Amphomycin destabilises the UptA:C_55_-P complex and sequesters C_55_-P, thus effectively out-competing the flippase for its carrier lipid substrate. (iv) UptA binds phospholipids, with a higher preference for the anionic than zwitterionic phospholipids. (v) UptA interacts with UptA more favourably than phospholipids, highlighting its preferred lipid substrate. (iv) Based on the proton-dependency of other DedA proteins (52) and the pH-dependency of UptA – C_55_-P interactions, we propose that the binding of H^+^ to UptA enables translocation of C_55_-P and its subsequent release for the next round of PG precursor synthesis.

Our studies also show that UptA selectively binds to C_55_-P over C_55_-PP and that the protein interacts more favourably with C_55_-P than the shorter chain analogues C_30_-P and C_20_-P. This implies that the hydrophobic tail of C_55_-P makes extensive contact with UptA, promoting the binding interactions and potentially facilitating the transport of carrier lipids. The higher affinity of UptA towards the monophosphate form of the carrier lipid, rather than to the diphosphate form, favours a mechanism in which UptA mediates the transport of C_55_-P rather than both C_55_-P and C_55_-PP. We further show that UptA interacts more favourably with the anionic lipids than the zwitterionic lipids and that these lipids are readily displaced by C_55_-P. This binding preference agrees with UptA being responsible for C_55_-P rather than the phospholipid translocation (23). Competition between anionic lipids, which are more abundant in the native lipid bilayer than C_55_-P, suggests that both lipids bind near the same sites on UptA and that the local lipid environment of UptA can potentially modulate its cellular function.

We provided molecular-level evidence that the conserved arginine (R112 and R118) residues in the putative membrane re-entrant helices of UptA are important for its substrate binding interactions, confirming earlier reports that *B. subtilis* strains bearing these mutations on UptA are more susceptible to inhibition by MX2401, a derivative of amphomycin (23). Despite the combined cellular pool of C_55_-PP and C_55_-P being limited to ∼1.5×10^5^ molecules per cell (55), they mediate the transport of glycan-bearing components of the cell wall peptidoglycan, teichoic acid and capsular polysaccharide (56). Therefore, antibiotics that block C_55_-P and its flippases are potential drug targets that can potentially inhibit multiple pathways and thus mitigate drug resistance. Accordingly, bacitracin and teixobactin form complex with C_55_-PP (57, 58) while amphomycin analogues inhibit bacterial cell wall formation by forming complexes with C_55_-P (23, 45, 59, 60). Herein we show that amphomycin and aspartocin D can additionally bind directly to UptA and induce dissociation of the UptA:C_55_-P complex, a finding that may guide the development of new antibiotic analogues to inhibit the flippase function of UptA and its homologues.

## MATERIALS AND METHODS

Detailed experimental procedures are provided in the SI Appendix. Briefly, UptA (wild type and mutants) was expressed in *E. coli* C43(DE3) cells (Lucigen) with GFP fusion on the C-terminus and solubilised from the membrane fraction by *n*-dodecyl-*β*-*D*-maltopyranoside (DDM). After purification by affinity chromatography, the GFP was cleaved by TEV protease, removed, and UptA was further purified by size-exclusion chromatography in a buffer containing 20 mM Tris-HCl (pH 8.0), 200 mM NaCl, 0.02% DDM. Before measurements, proteins were buffer-exchanged into 0.05% LDAO, 200 mM ammonium acetate at the desired pH using a centrifugal buffer exchange device (Micro Bio-Spin 6, Bio-Rad). Stock solutions of lipids were prepared from chloroform/methanol solution by evaporating aliquots of known volume in a SpeedVac. After drying, the lipid films were weighed and resuspended to a final concentration of 0.5 mM in 200 mM ammonium acetate and 0.05% LDAO by vertexing. Stock solutions of 0.5 mM amphomycin, aspartocin D, daptomycin, polymyxin B, polymyxin E, and bacitracin were made in the same buffer. All experiments were repeated three times from newly prepared stock solutions. Native mass spectrometry was performed on a Q-Exactive hybrid quadrupole-Orbitrap mass spectrometer (Thermo Fisher Scientific, Bremen, Germany) and lipid identification was performed on an Orbitrap Eclipse Tribrid mass spectrometer (Thermo).

## ACKNOWLEDGMENTS

Research in the C.V.R. laboratory is supported by a Wellcome Trust Award (221795/Z/20/Z); and a Medical Research Council (MRC) programme grant (MR/V028839/1) on which A.O.O. is a researcher co-investigator. J.R.B. acknowledges the Royal Society University Research Fellowship grant (URF\R1\211567). A.O.O. is a Junior Research Fellow at St Anne’s College, University of Oxford, United Kingdom.

## Supporting Information

### Methods

#### Chemicals

C_55_-PP, C_55_-P, C_30_-P and C_20_-P were purchased from Larodan. Phospholipids – a15:0-i15:0 PE, a15:0-i15:0 PG, 16:0-18:1 PE, 16:0-18:1 PG, 18:1-18:1 PE, 18:1-18:1 PG – were from Avanti Polar Lipids. Amphomycin and aspartocin D were purchased from Cayman; while Daptomycin and bacitracin were purchased from Merck. *n*-dodecyl-*β*-D-maltoside (DDM), *n*-decyl-*β*-D-maltoside (DDM), n-octylglucoside (OG) and *n*-dodecyl-*N,N*-dimethylamine-*N*-oxide (LDAO) were from Anatrace.

#### Plasmids

A codon-optimised version of *Bacillus subtilis* UptA (yngC) (Integrated DNA Technologies) was inserted into a modified pET15b vector between NdeI and NheI endonuclease restriction sites by In-Fusion HD cloning kit (Takara Bio). The resulting plasmid encodes UptA fused to GFP-His_6_ on the C-terminus via a TEV-protease cleavable site. Plasmids for UptA mutants were generated by site-directed mutagenesis with CloneAmp HiFi PCR Premix (Takara Bio) using pairs of oligonucleotides listed in Table S1. The plasmid pET26-MBP-MraYaa bearing HRV3C protease cleavage site between the MBP and the wild-type MraY was a kind gift from S-Y Lee (Addgene plasmid #100166) (1).

**Table S1:**
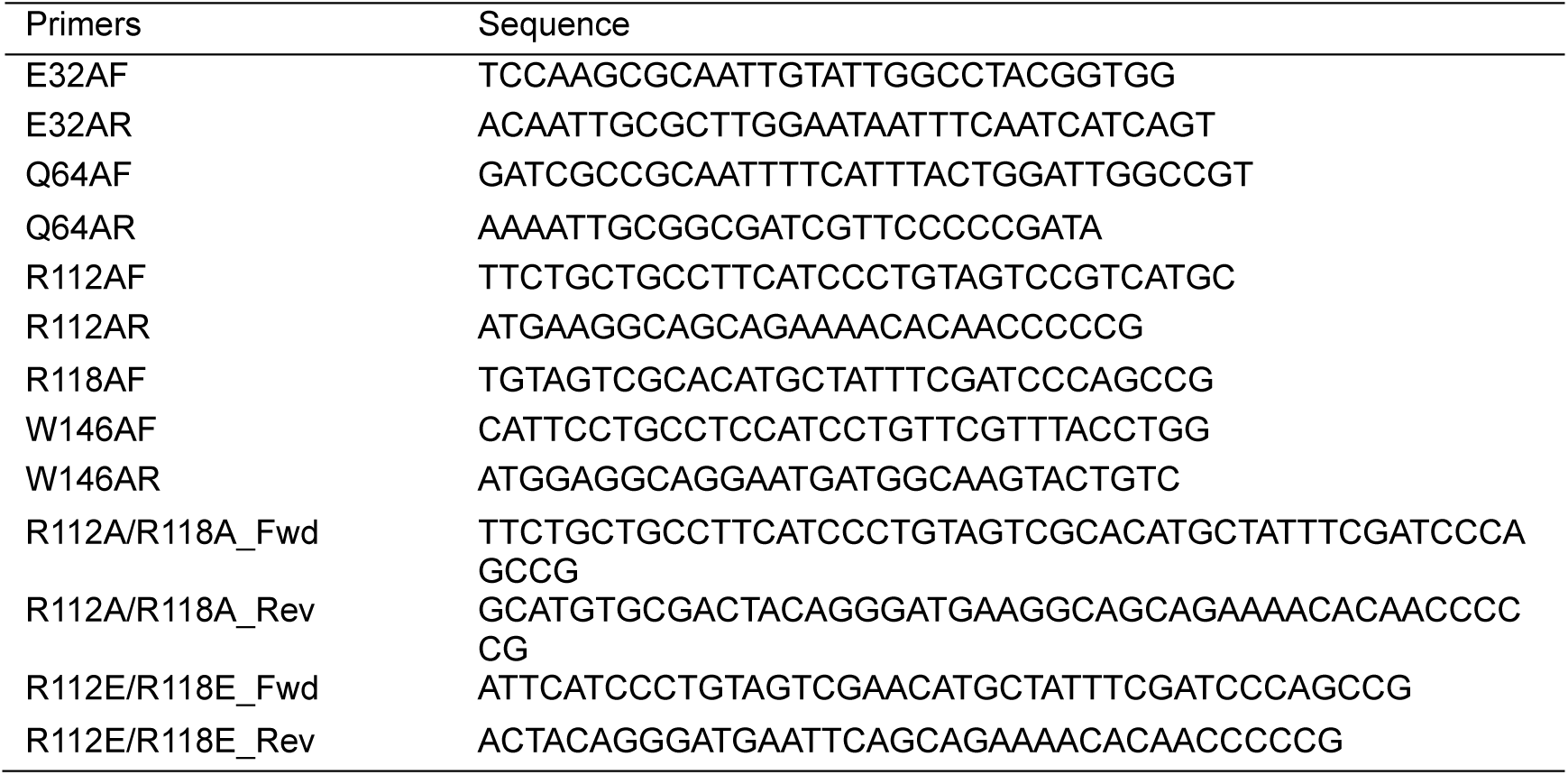
Oligonucleotides used to generate plasmids for the expression of UptA mutants.

#### Expression and purification of UptA

Plasmids were used to transform chemically competent *E. coli* C43(DE3) cells (Lucigen) and selected on LB/Agar containing 100 μg/ml ampicillin at 37 °C. Single colonies were used to inoculate 60 mL LB containing 100 μg/ml ampicillin and were grown at 37 °C overnight for 16 h. For large-scale protein expression, 10 mL of the overnight culture was aseptically transferred into 1 L of LB/ampicillin and cells were allowed to grow at 37°C until the OD_600 nm_ reached 0.6-0.8 after which the temperature was lowered to 18 °C. Expression of UptA-GFP was induced by adding isopropyl-β-D-thiogalactopyranoside (IPTG) at a final concentration of 0.5 mM UptA-GFP and shaking for 20 h at 18 °C. Cells were harvested by centrifugation at 5000 × *g* and lysed immediately or stored at -80 ° C until required.

Cells were thawed on ice and resuspended in a buffer containing 50 mM Tris-HCl (pH 7.4), 200 mM NaCl, 10 µg/mL deoxyribonuclease I (DNAse I), 5 mM MgCl_2_, 5 mM β-mercaptoethanol and EDTA-free protease inhibitor cocktail tablets. The cells were then disrupted with a microfluidiser (Microfluidics) operated at 20,000 psi. Non-lysed cells and debris were pelleted by centrifugation at 20,000 × *g* for 20 min and the clarified lysate was retained. The membranes fraction was then pelleted by ultracentrifugation at 100,000 x g for 1 h. They were resuspended in a buffer containing 20 mM Tris-HCl (pH 8.0), 300 mM NaCl, 20% glycerol. Membrane aliquots were either solubilised immediately or flash-frozen in liquid nitrogen and stored at -80°C. The membrane was solubilised by incubation with DDM at a final concentration of 1% (w/v) for 60 min. Non-solubilised aggregates were removed by centrifugation at 20,000×g for 20 min and the supernatant was loaded onto a 5-ml HisTrap HP column preequilibrated in buffer A (20 mM Tris-HCl (pH 8.0), 200 mM NaCl, 10% glycerol, 0.03% DDM and 20 mM imidazole). After binding of the His-tagged protein, the column was washed with 100 mL of buffer A, and then with 50 mL of buffer B (20 mM Tris-HCl (pH 8.0), 200 mM NaCl, 10% glycerol, 0.03% DDM and 50 mM imidazole). UptA-GFP was eluted from the column with buffer C (20 mM Tris-HCl (pH 8.0), 200 mM NaCl, 10% glycerol, 0.02% DDM and 300 mM imidazole). To remove imidazole and to cleave the GFP fusion, eluted protein was mixed with TEV protease and dialysed against buffer D (20 mM Tris-HCl (pH 8.0), 150 mM NaCl, 10% glycerol, 0.02% DDM) at 4°C for 16 h.

The digestion mixture was subsequently incubated with 2-mL bed volume of nickel nitriloacetic acid (NTA) agarose resin to remove His-tagged GFP and TEV protease. Tag-free UptA was collected as the flowthrough, concentrated using an Amicon centrifugal concentrator with nominal molecular weight cut-off 50 kDa. The protein was finally purified by size-exclusion chromatography (SEC) using a 24-mL Superdex 200 Increase column. The SEC buffer was 20 mM Tris-HCl (pH 8.0), 200 mM NaCl, 0.02% DDM. UptA was concentrated to ∼200 uM, aliquoted, flash-frozen flash in liquid nitrogen, and stored at -80 °C until use. Concentration was measured using a Biomate UV spectrophotometer at 280 nm using extinction coefficients calculated from their predicted amino acid sequences. UptA mutants were purified in 3 biological replicates, using the same procedures as for the wild type.

#### Expression and purification of MraY

MraY from *Aquifex aeolicus* was purified as previously described (3). Briefly, the plasmid pET26-HisMBP-MraYaa was used to transform chemically competent *E. coli* C41(DE3) cells (Lucigen) and selected on LB/Agar containing 50 μg/ml kanamycin at 37 °C. Cells were grown in Terrific Broth media supplemented with 50 μg/ml kanamycin at 37°C until the OD_600 nm_ reached 0.8. Protein expression was induced by adding IPTG at a final concentration of 1 mM for 4 h at 37 °C. Cells were harvested by centrifugation at 5000 × *g* and lysed and solubilized by incubation with 1% DDM for 1 h. His-tagged MBP-MraY was purified by affinity chromatography and digested by HRV3C protease. MraY was isolated by SEC into a buffer containing 20 mM Tris-HCl (pH 8.0), 200 mM NaCl, 0.02% DDM.

#### Denaturing SDS-PAGE

Denaturing SDS-PAGE was performed using a precast NuPAGE™ Novex 12% Bis-Tris Mini protein gel (Invitrogen) according to the manufacturer’s specifications. 30 μL of 0.3 μg/μL protein were mixed with 10 μL of 4X LDS sample loading buffer (Invitrogen) and heated at 80 °C for 5 min. 20 μL of the resulting samples were then loaded into in each well. 5 μL PageRuler™ prestained protein ladder (Thermo Scientific) was loaded into a well as molecular weight marker. Electrophoresis was performed at room temperature for 36 min using a constant voltage (200 V) in 1X solution of NuPAGE™ MES SDS running buffer (Invitrogen). The gel was stained with Quick Coomassie stain solution (Neo Biotech) for 60 min and rinsed with water for 24 h before imaging on iBright™ FL1500 imaging system (Thermo).

Lipid extracts from *B. Subtilis*

*B. subtilis* strain 168 (ATCC 23857) was cultivated in 200 mM LB media at 37 °C until OD_600 nm_ 1.2 under aurobic conditions. Cells were harvested by centrifugation at 4,000 *g* for 20 minutes and resuspended in 20 mM Tris-HCl (pH 8.0), 150 mM NaCl. Cells were lysed by 5-6 passes through a microfluidizer at 20,000 psi. The lysates were clarified by centrifugation at 4,000 *g* for 10 minutes, and the totoal membrane was pelleted at 120,000 *g* for 1 h. The membrane pellet was ressupended in 20% sucrose, 50 mM Tris-HCl (pH 8.0), and loaded as the topmost layer of a 4-layer sucrose gradient: 20% (4 mL), 36% (3 mL), 51% (8 mL) and 55% (2 mL) (4). After centrifugation at 120,000 *g* for 20 hrs, the cytosolic membrane was collected at the interface of 36% and 45% sucrose layers, and washed 3 times with 1 M ammonium acetate by dilution-centrigugation at 120,000 *g* for 30 min. The membrane pellet was homogenised in 2 mL of 1 M ammonicum acetate and stored in -80 °C until use. Lipids were extracted by homogenising 0.8 mL of membrane with 1 mL of 1:1 vol/vol) chloroform-methanol mixture in a glass vial followed by vortexing. To separate the aqueous and the organic pahses, adding 1 mL chloroform was added, and the mixture was centrifuged at 3,000 *g* for 5 min. The chloroform layer containing the total lipids was recovered into a clean glass vial, and the organic solvent was evaporated in a SpeedVac for 16 h. Lipid film was resuspended 200 mM ammonium acetate, 0.05% LDAO (pH 8.0) to ∼0.5 mg/mL for the binding experiments.

#### Sample preparation for native MS

Before measurements, the protein was buffer-exchanged into 0.05% LDAO, 200 mM ammonium acetate (“MS buffer”) at the desired pH using a centrifugal buffer exchange device (Micro Bio-Spin 6, Bio-Rad). Stock solutions of lipids were prepared from chloroform/methanol solution by evaporating aliquots of known volume in a SpeedVac. After drying, the lipid films were weighed and resuspended to a final concentration of 0.5 mM in 200 mM ammonium acetate (pH8) and 0.05% LDAO by vertexing. Amphomycin and aspartocin D were purchased from Cayman while Daptomycin and bacitracin were purchased from Merck. Stock solutions of 0.5 mM amphomycin, aspartocin D, daptomycin and bacitracin, were made in the same buffer. All experiments were repeated three times from newly prepared stock solutions.

#### Native mass spectrometry

About 3 μL protein aliquot was transferred into a gold-coated borosilicate capillary (Harvard Apparatus) and was mounted on the nano ESI source of a Q-Exactive hybrid quadrupole-Orbitrap mass spectrometer (Thermo Fisher Scientific, Bremen, Germany). The instrument settings were 1.2 kV capillary voltage, S-lens RF 200%, argon UHV pressure 3.3 × 10^−10^ mbar, the capillary temperature was set to 200 °C, and resolution of the instrument was set to 17,500 at a transient time of 64 ms. Voltages of the ion transfer optics –injection flatapole, inter-flatapole lens, bent flatapole, and transfer multipole were set to 5, 3, 2, and 30 V respectively. The noise level was set at 3. Unless otherwise stated, proteins were activated by applying 75 V in the high-energy collisional dissociation cell without in-source trapping. For the MS^2^ spectra, parent ions were isolated using an in-source trapping voltage of -200 V, and then fragmented by applying 150-200 V in the high-energy collision-induced dissociation (HCD) cell. Lipids bound to UptA from the total lipid extracts were identified on an Orbitrap Eclipse Tribrid mass spectrometer (Thermo). Peaks corresponding to the parent ions in the negative ESI spectra were isolated within ±1.0 Th and then activated with HCD voltage 10-30%. To identify endogenous modifications using native top-down mass spectrometry, the 6+ charge state of monomeric UptA was isolated (3828 ± 5 Th) and activated using HCD (200 V) with reduced trapping pressures. The resulting fragments were detected in the Orbitrap (R = 280,000 @ *m*/*z* 200), averaging ∼500 transients to maximise the signal-to-noise ratio. Theoretical fragments from N-terminally formylated UptA were matched to the MS^2^ spectrum (3 ppm) using TDValidator (Proteinaceous) and manual validation. Data were visualised and exported for processing using the Qual browser of Xcalibur 4.1.31.9 (Thermo Scientific).

Spectral deconvolution was performed using UniDec 6.0.2 (2). Relative binding affinities were obtained from deconvoluted spectra by dividing the intensity of ligand-bound protein peaks by the sum of the intensities of ligand-bound and ligand-free protein peaks. The mean and standard deviation of these fractional binding intensities from three independent experiments were plotted against the concentrations of the ligand C_55_-P or C_55_-PP. To obtain the apparent dissociation constant *K*_d_, the mean relative intensities at different concentrations were fitted globally using GraphPad Prism 10.2.0 to Eq. (1):

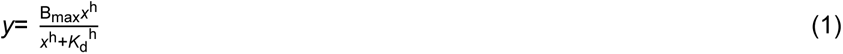

Where y is the mean relative intensity and x is the corresponding ligand concentration. B_max_ is the maximum specific binding, *K*_d_ is the apparent dissociation constant, and h is the Hill coefficient.

## Supplementary Figures

**Fig. S1.**
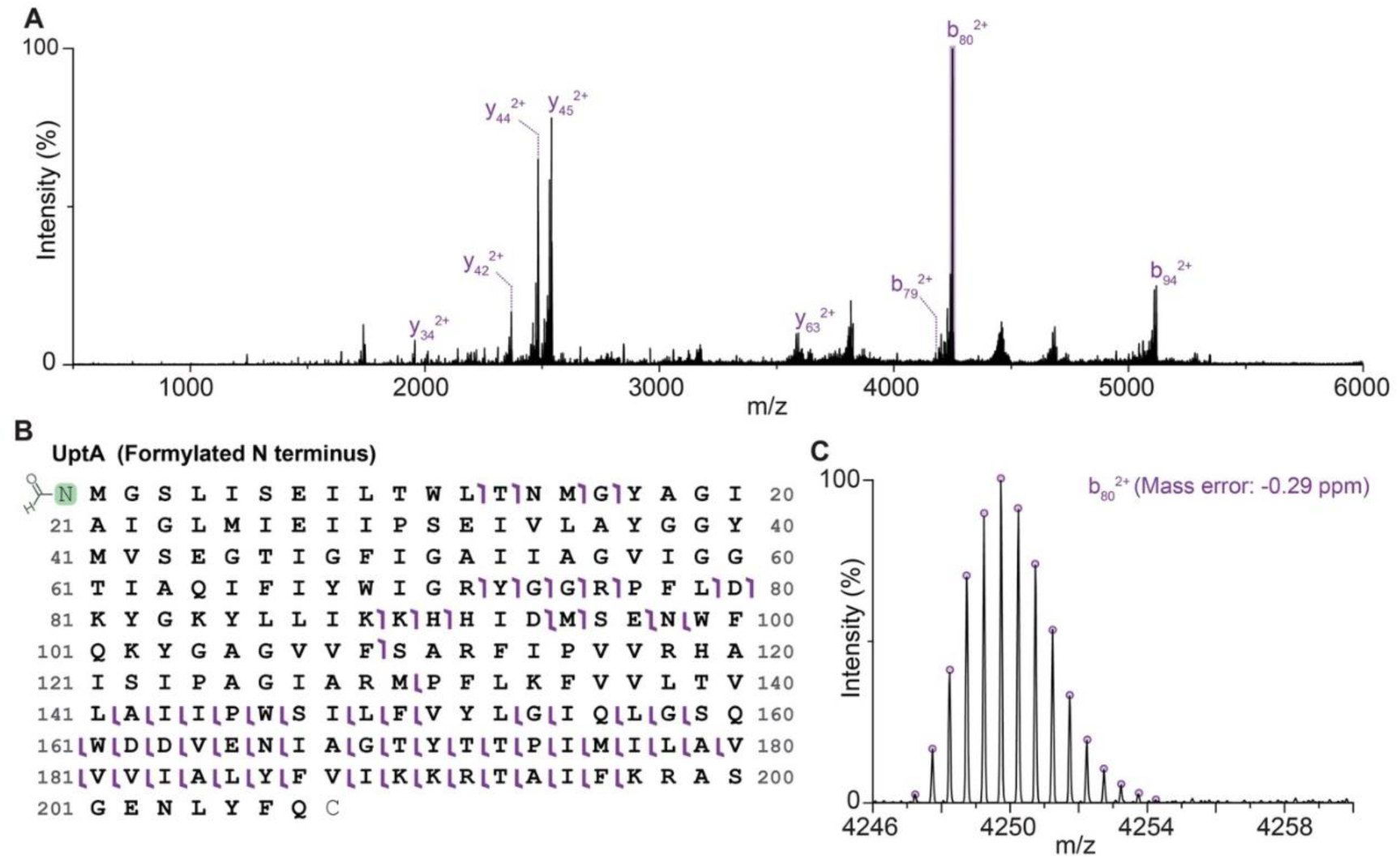
UptA is formylated at the N-terminus. **A.** HCD MS/MS spectrum of UptA (6+). The eight most abundant assigned ions are labelled. **B**. Sequence map of the UptA construct used in this study. Terminal *b*- and *y*-type fragment ions observed in the MS/MS spectrum are indicated. **C.** Expanded view of the peak corresponding to b_80_^2+^ ion. The theoretical isotopic envelope is overlaid.

**Fig. S2.**
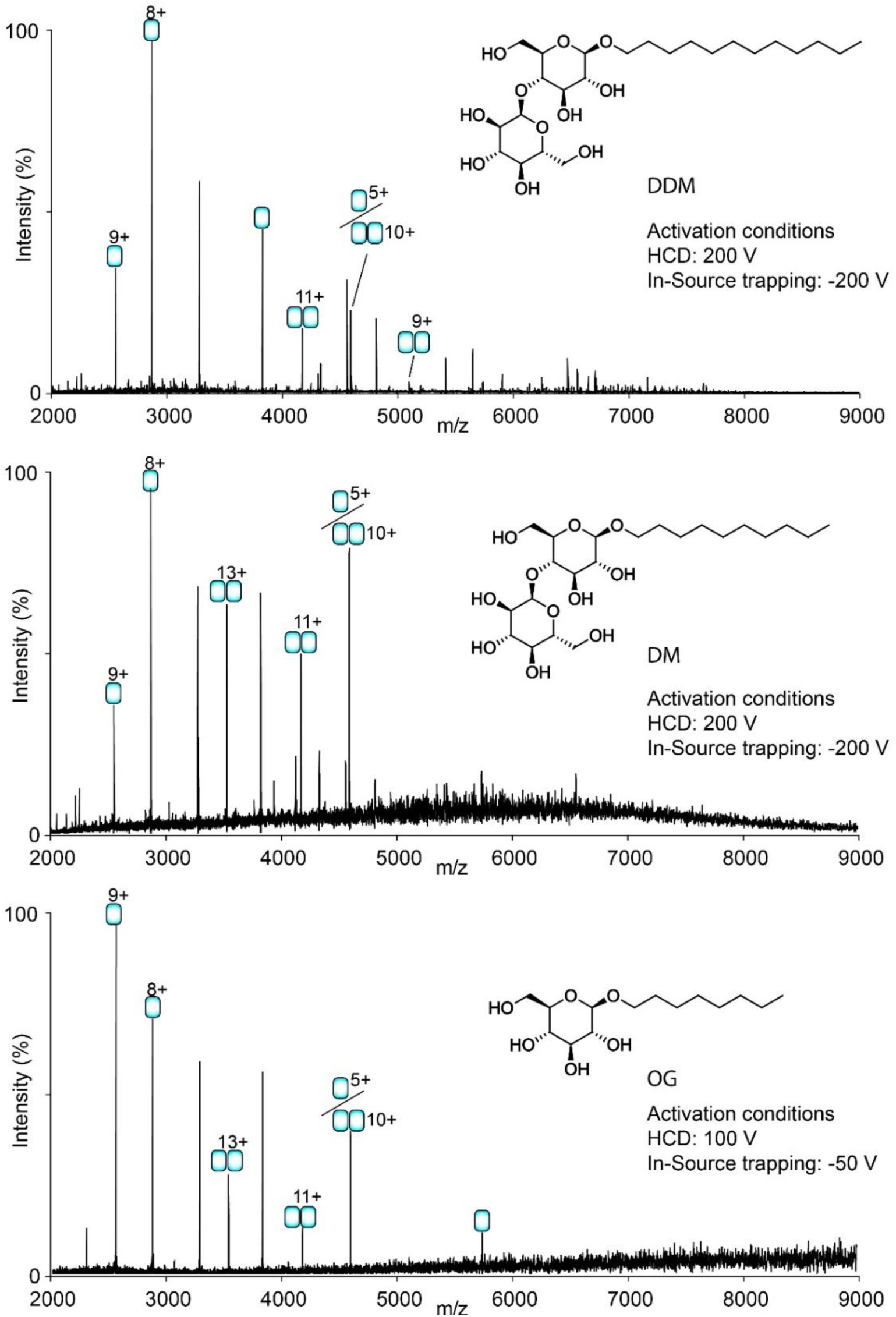
Oligomeric state of UptA in different detergents. Shown are the spectra recorded for UptA in buffer containing 200 mM Ammonium acetate (pH 8.0) supplemented with indicated detergents. The instrument was tuned differently in each case to maximise the transmission of dimeric species.

**Fig. S3.**
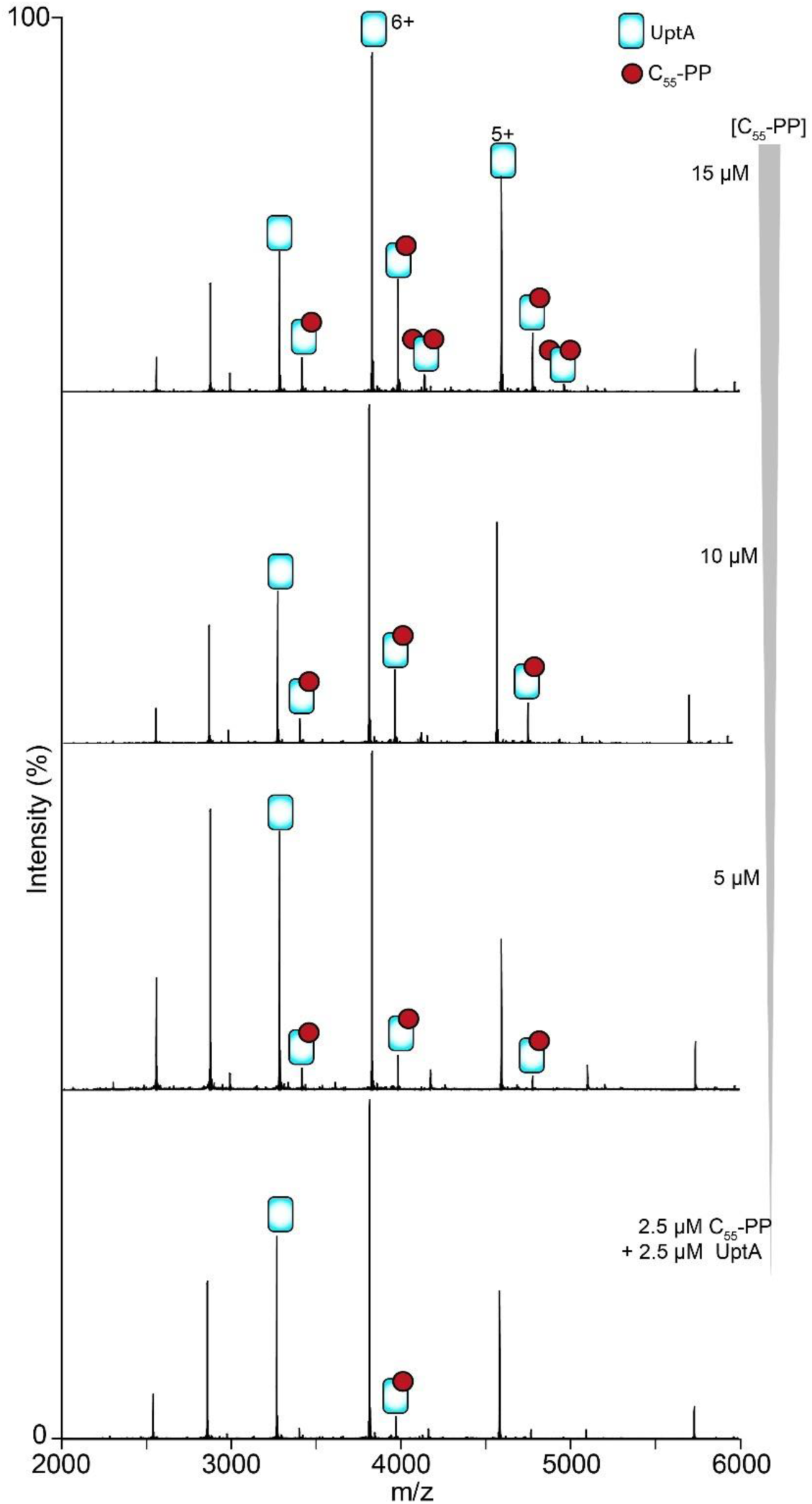
UptA:C_55_-PP binding interactions. Spectra for UptA incubated with different concentrations of C_55_-PP in a buffer containing 200 mM ammonium acetate (pH 8), 0.05% LDAO. UptA bind C_55_-PP but with lesser intensities compared to C_55_-P (cf. Fig. 2A).

**Fig. S4.**
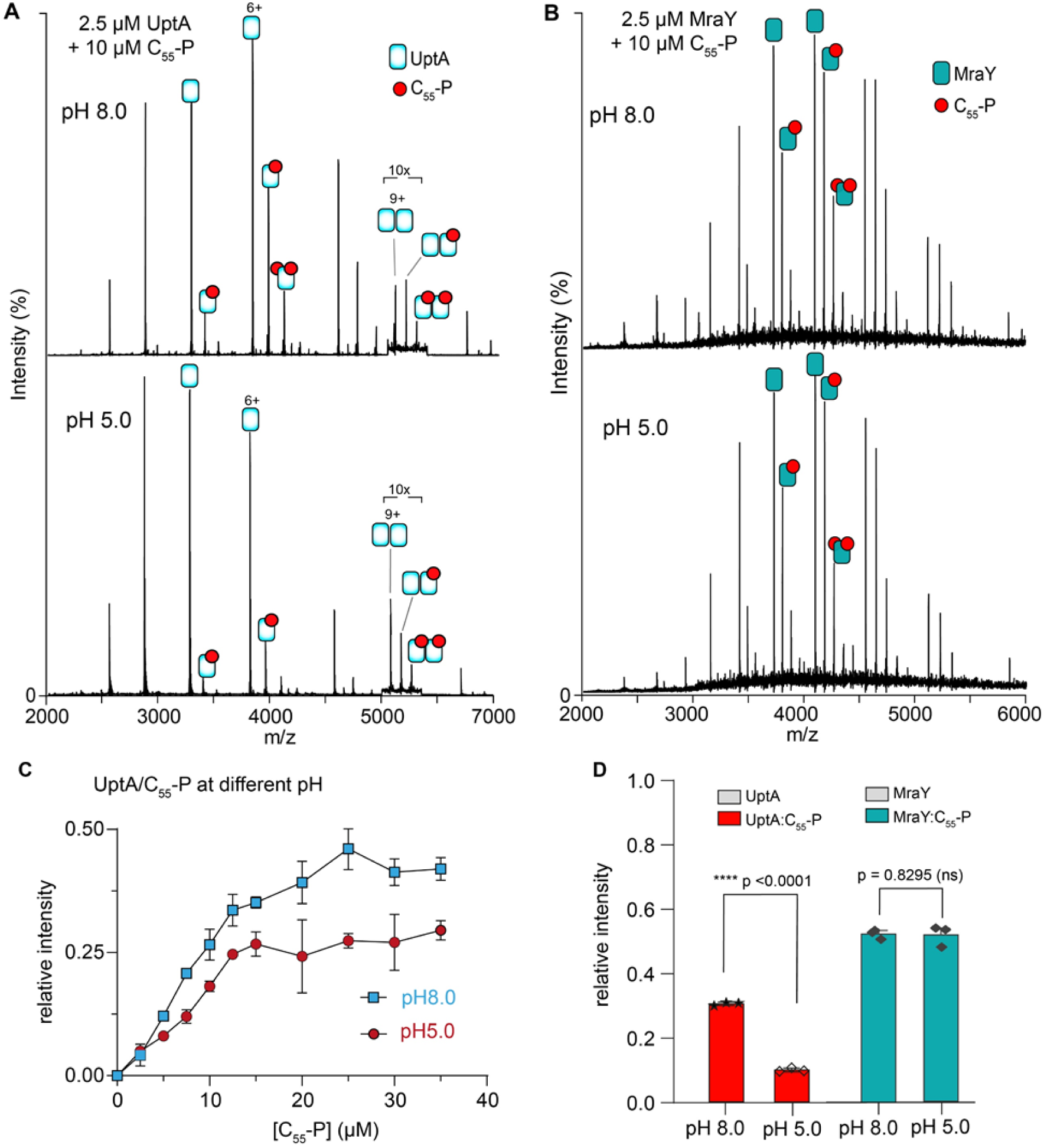
pH dependency of C_55_-P binding interactions with UptA and MraY. **A,B**. Spectra for UptA/C_55_-P and MraY/C_55_-P mixtures at pH 8.0 and pH 5.0. Peaks corresponding to the dimeric form of UptA is observed mostly in complex with C_55_-P. **C.** Relative intensity of UptA:C_55_-P at pH 8.0 and pH 5.0 as a function of concentrations of C_55_-P in a buffer containing 200 mM ammonium acetate,0.05% LDAO. Data point is an average of 3 independent replicates, and the error bars are standard deviations. **D.** Relative intensity of C_55_-P (10 µM) bound to UptA (2.5 µM) and MraY (2.5 µM) at different pH. Unlike UptA, there is no pH-dependency in C_55_-P binding by MraY. Bar represents the mean of 3 replicate measurements shown as data points, and the error bars are standard deviations.

**Fig. S5.**
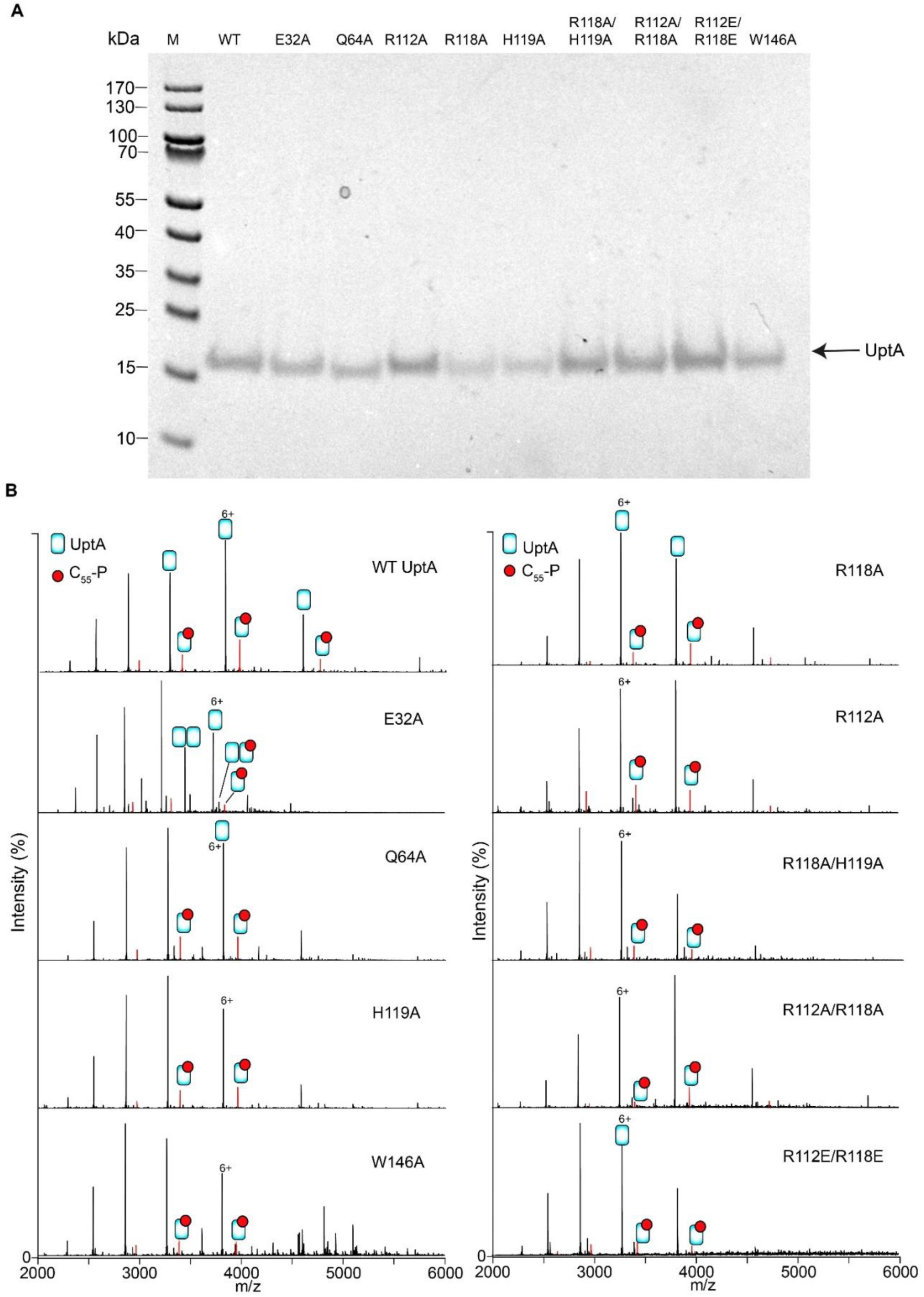
Native MS analysis of C_55_-P binding to UptA mutants. **A.** SDS-PAGE image of UptA wild-type (WT) and mutants. M, PageRuler^TM^ prestained protein ladder. UptA is ∼22 kDa but migrate on SDS-PAGE as ∼17 kDa protein. **B.** Representative spectra for UptA wild type and mutants (5 µM) incubated with C_55_-P (10 µM). Proteins were liberated from a buffer containing 0.05% LDAO and 200 mM ammonium acetate (pH 8.02) using a collisional activation of 100 V.

**Fig. S6.**
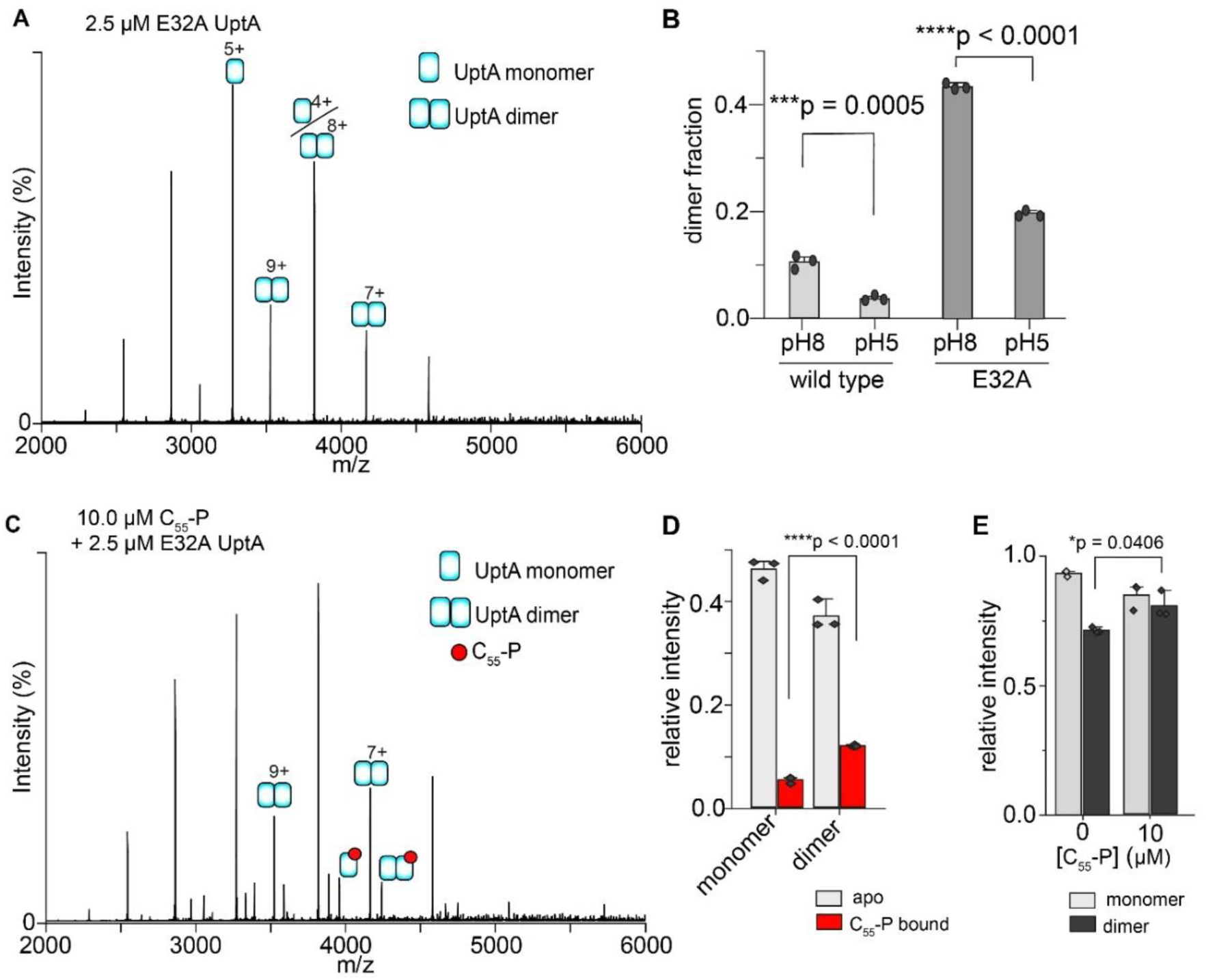
Dimer fraction of UptA binds C_55_-P more than the monomeric form. **A.** Spectrum of UptA E32A mutant display peaks corresponding to monomers and dimers. **B.** Relative intensity of dimer populations in spectra for the wild-type UptA and the E32A mutant. In both cases, the dimer fraction was higher at pH 8.0 than at pH 5.0. **C.** Spectrum of UptA E32A mutant equilibrated with 10 µM C_55_-P. **D.** Corresponding intensities of apo and C_55_-P bound UptA. Dimeric form of UptA bind to the ligand C_55_-P more significantly than the monomeric form. **E.** The relative proportion of UptA monomers and dimers in a mixture of 2.5 µM E32A UptA and 10 µM C_55_-P at pH 8.0. The ligand C_55_-P caused a modest increase in the dimer fraction.

**Fig. S7.**
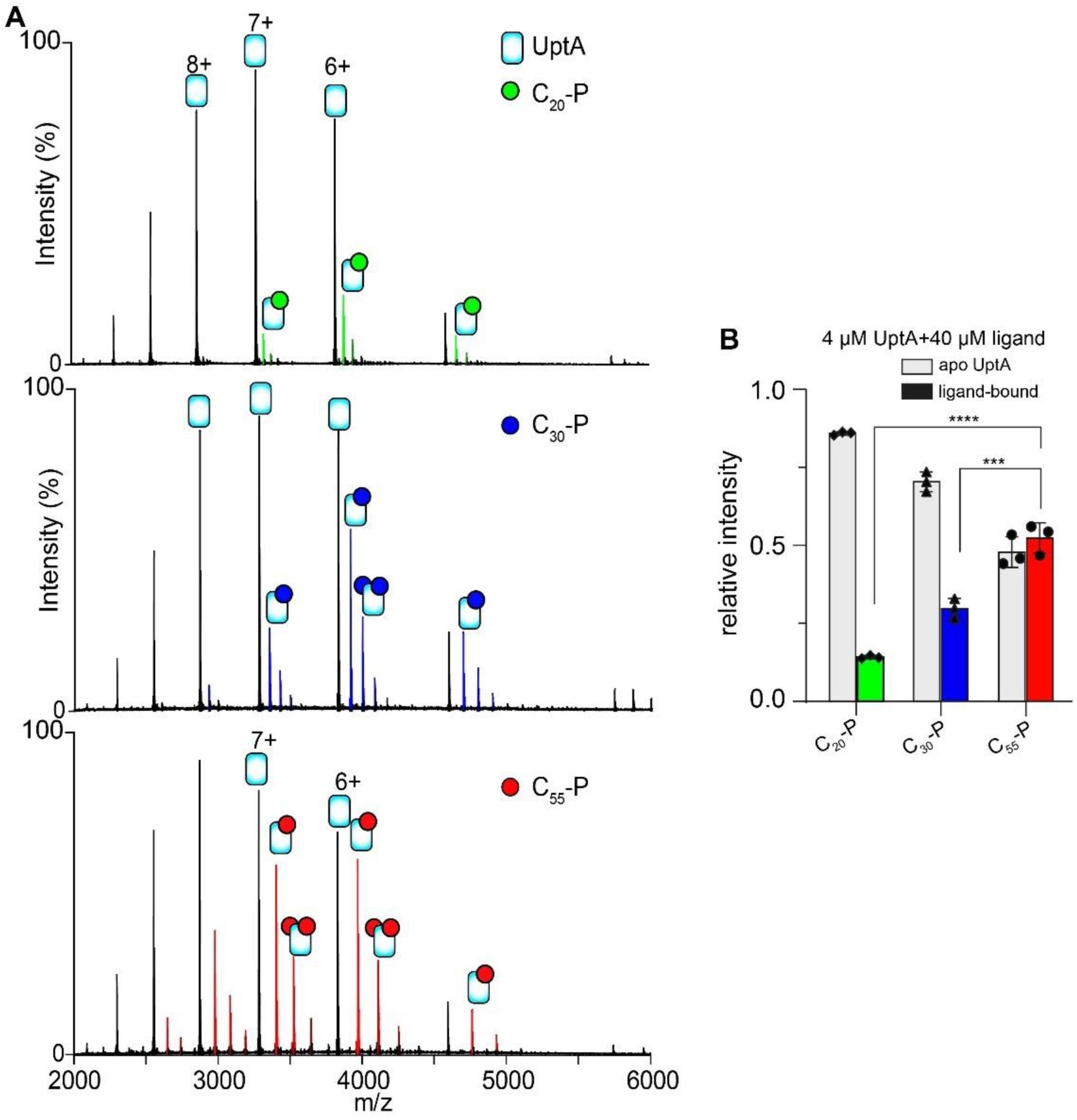
UptA binds carrier lipids of longer chain lengths more intensely than those of shorter chain, and phospholipids. **A.** Spectra for 4 µM UptA equilibrated with 40 µM undecaprenyl phosphate, C_55_-P hexaprenyl phosphate, C_30_-P and geranylgeranyl phosphate, C_20_-P. Samples were prepared in a buffer containing 200 MM ammonium acetate (pH8.0) and 0.05% LDAO. **B.** Relative intensity of ligand-free and ligand-bound UptA. UptA binds to C_55_-P more intensely than the shorter chain analogues.

**Fig. S8.**
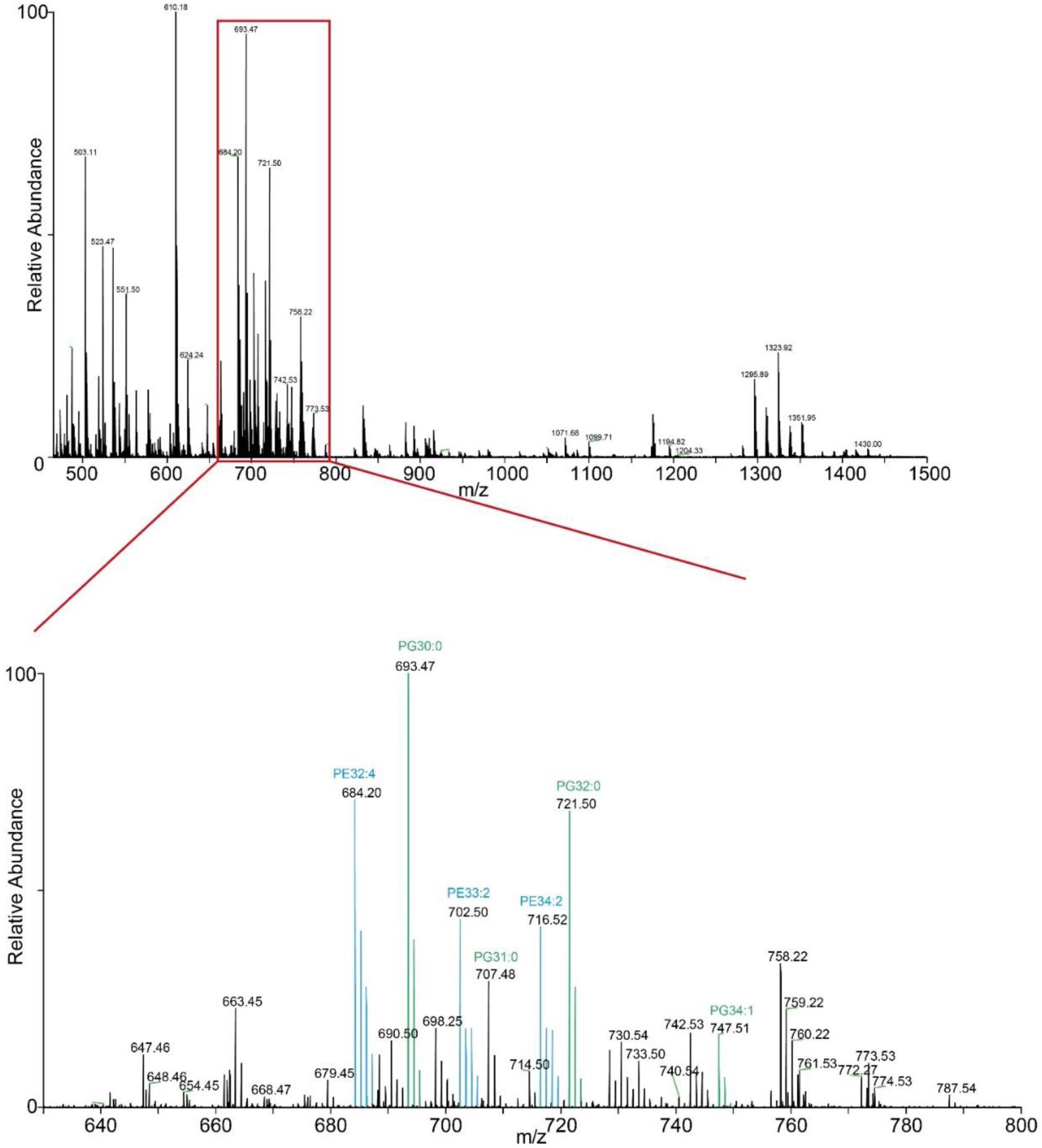
Mass spectrum of total lipid extract from B. subtilis membrane. Lipid extracts from B. subtilis membrane was dissolved in 0.2 M ammonium acetate, 0.05% LDAO (pH8.0) and analysed in the positive ESI mode using source activation of 25 V. Main phospholipids (PE, blue; PG, green) are highlighted in the expanded view.

**Fig. S9.**
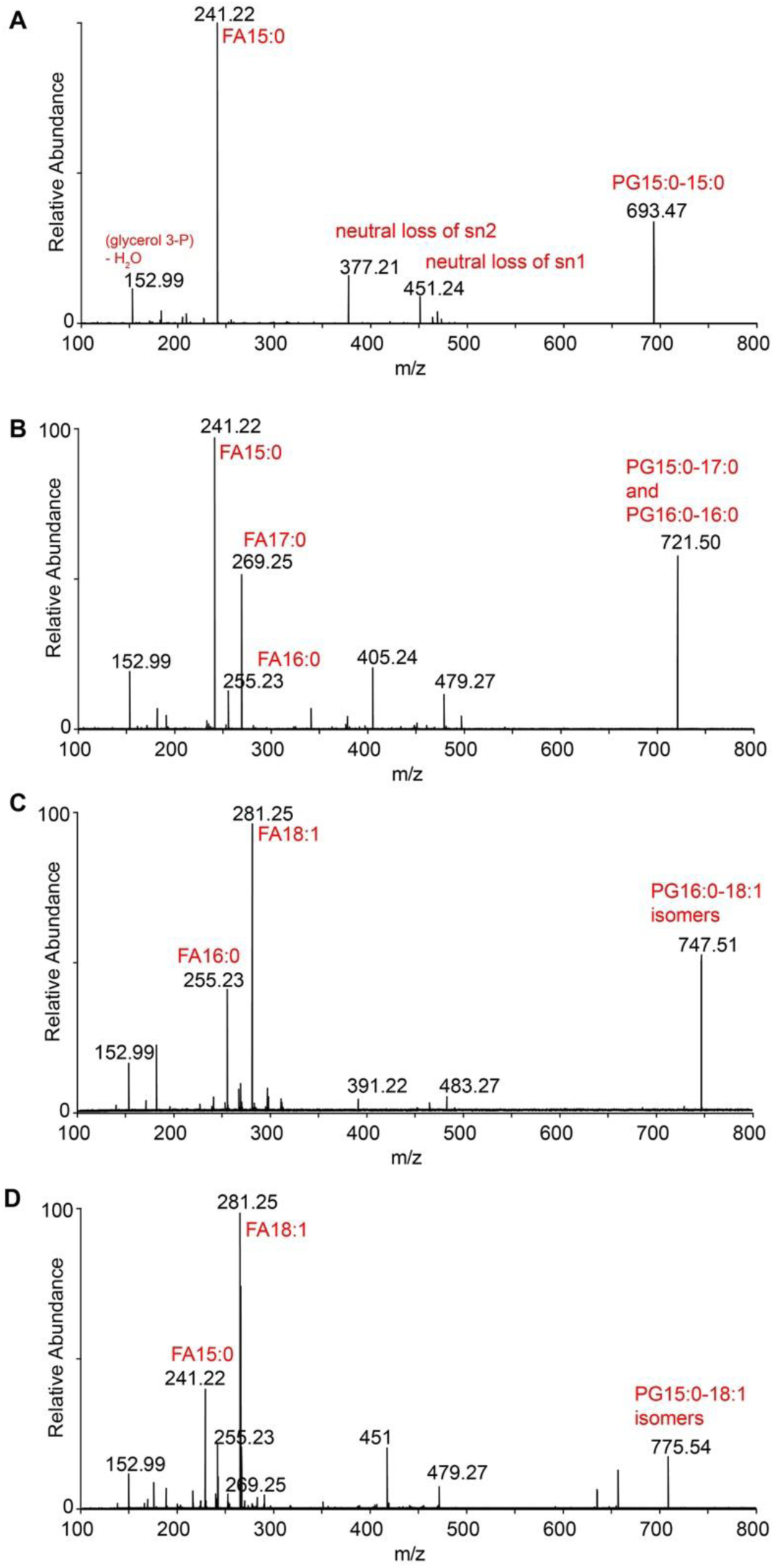
MS/MS spectra (negative ESI mode) for lipid species interacting with UptA from the total lipid extracts of *B. subtilis*. Peaks corresponding by mass to each of the parent ions were isolated with a source activation of 150 V, and then fragmented with 15-25 V in the HCD cell. All species carry a single negative charged.

**Fig. S10.**
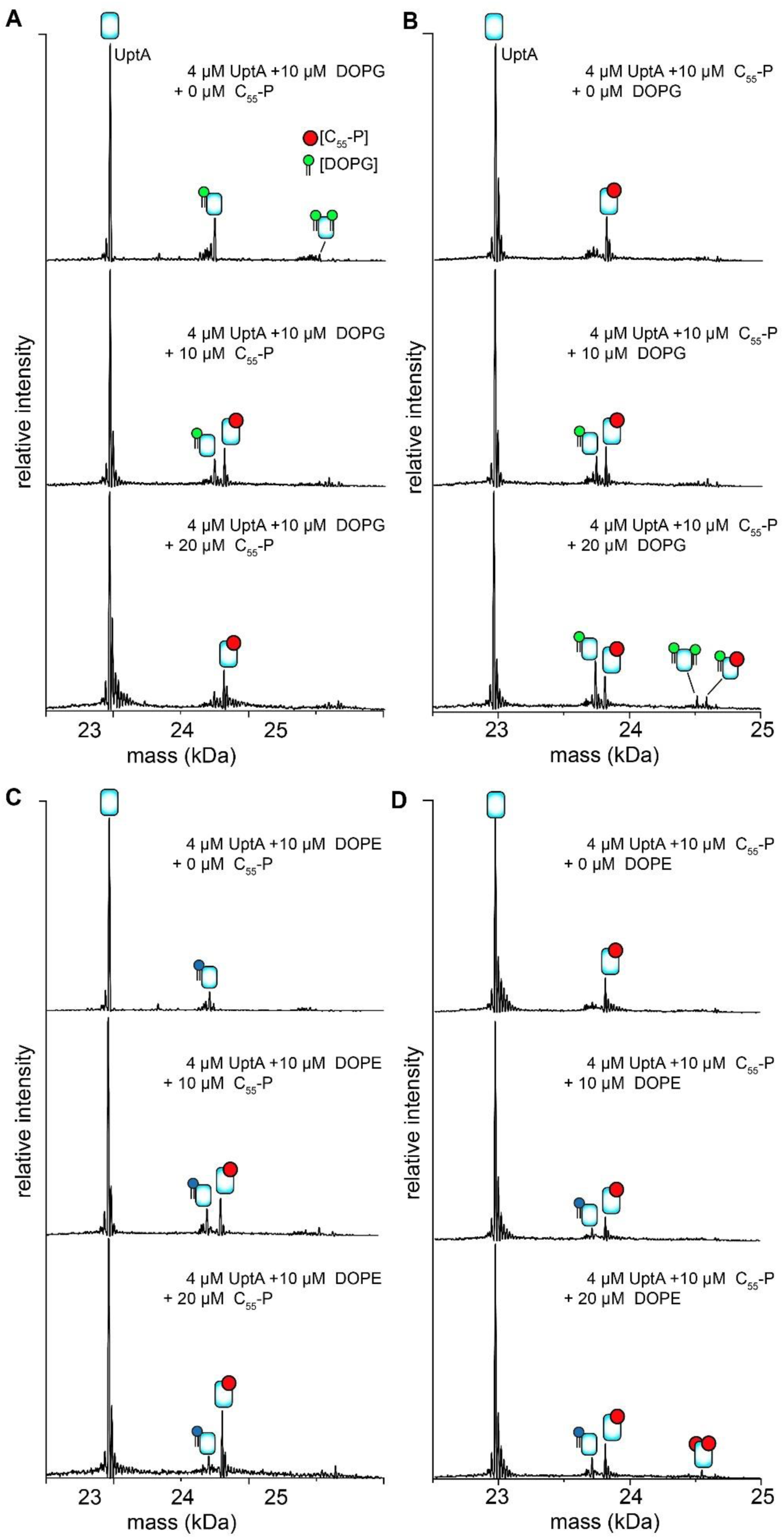
C55-P outcompetes phospholipids for UptA binding. **A,B**) Deconvoluted native mass spectra for 4 µM UptA and10 µM 18:1-18:1 PG (DOPG, panel A) or 18:1-18:1 PE (DOPE, panel B) in the presence of increasing concentrations of C_55_-P. The intensities of peaks assigned to lipid bound to UptA attenuates upon titration with C_55_-P. **C,D**) Spectra for 4 µM UptA and 10 µM C_55_-P, then incubated with increasing concentrations of DOPG (Panel B) or DOPE (panel (D). The peaks assigned to UptA-bound C_55_-P remained in the presence of lipids, indicating that C_55_-P binds to UptA more favourable than the phospholipids.

**Fig. S11.**
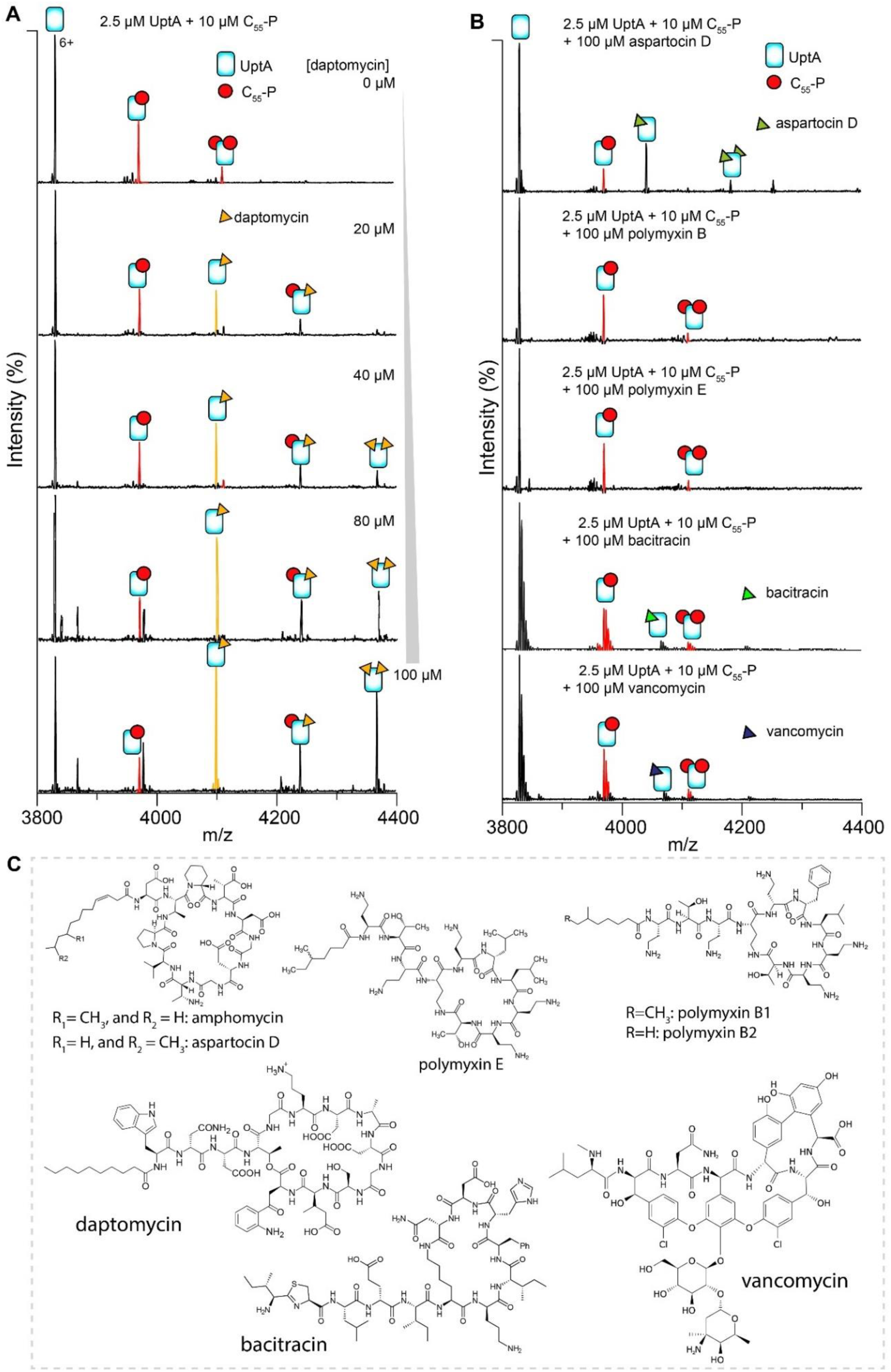
Impact of cell-wall antibiotics on UptA:C_55_-P interactions. **A.** Spectra (6+ charge state) of 2.5 µM UptA and 10 µM C_55_-P in the presence of different daptomycin concentrations. Unannotated peaks correspond to multiple daptomycin binding events in the 5+ charge state. **B**. Spectra (6+ charge state) of UptA and C_55_-P in the presence of different antibiotics and acquired using the same instrument settings and collisional activation of 75 V. Relative intensity of UptA-bound C_55_-P in the spectra shown in Figure 5. **C**. Chemical structures of lipopeptide antibiotics used in this study.

